# Global cerebrospinal fluid circulation mapping using gold nanoparticle enhanced X-ray microtomography reveals region-specific brain and spinal cord CSF pathways

**DOI:** 10.1101/2021.12.18.473250

**Authors:** Shelei Pan, Dakota DeFreitas, Sruthi Ramagiri, Peter Bayguinov, Carl D. Hacker, Abraham Z. Snyder, Jackson Wilborn, Hengbo Huang, Peter H. Yang, Dhvanii K. Raval, Sanja Svben, Samuel Achilefu, Rui Tang, Gabriel Haller, James D. Quirk, James A. Fitzpatrick, Prabagaran Esakky, Jennifer M. Strahle

**Affiliations:** Department of Neurosurgery, Washington University School of Medicine, Washington University in St. Louis, St. Louis, MO 63110, USA; Washington University Center for Cellular Imaging, Washington University School of Medicine, Washington University in St. Louis, St. Louis, MO 63110, USA.; . Department of Radiology, Washington University School of Medicine, Washington University in St. Louis, St. Louis, MO 63110, USA.; . Department of Biomedical Engineering, Washington University School of Medicine, Washington University in St. Louis, St. Louis, MO 63110, USA.; . Department of Genetics, Washington University School of Medicine, Washington University in St. Louis, St. Louis, MO 63110, USA; . Departments of Neuroscience and Cell Biology and Physiology, Washington University School of Medicine, Washington University in St. Louis, St. Louis, MO 63110, USA; . Department of Orthopedic Surgery, Washington University School of Medicine, Washington University in St. Louis, St. Louis, MO 63110, USA; . Department of Pediatrics, Washington University School of Medicine, Washington University in St. Louis, St. Louis, MO 63110, USA

## Abstract

Cerebrospinal fluid (CSF) movement within the brain interstitium is essential for the development and functioning of the brain. However, the interstitium has largely been thought of as a single entity through which CSF circulates, and it is not known whether specific cell populations within the CNS preferentially interact with CSF. Here, we developed a novel technique for CSF tracking, gold nanoparticle enhanced X-ray microtomography, to achieve micrometer-scale resolution visualization of CSF pathways during development. Using this method and subsequent histological analysis, we map global CSF pathways and present novel particle size-dependent circulation patterns through the CNS. We identify an intraparenchymal CSF circulation that targets stem cell-rich and cholinergic neuronal populations. CSF solute distribution to these areas is mediated by CSF flow along projections from the basal cisterns which is altered in posthemorrhagic hydrocephalus. Our study uncovers region-specific patterns in a biologically driven CSF circulation that has implications for normal brain development and the pathophysiology of hydrocephalus and neurodegenerative disorders.

## Introduction

Postnatal brain development involves dramatic changes in the cells and structure of the brain and the establishment of interconnecting signaling pathways.^1^ Numerous studies have implicated roles for the environment around developing neurons, including the mature neurons and glia, growth factors, cytokines, interstitial fluid (ISF), and cerebrospinal fluid (CSF), as playing key roles in neural development and activity.^2, 3^ Many neural precursor populations, including the subventricular zone and hippocampus, are in direct contact with the CSF in the brain ventricles and are thought to be modulated by CSF constituents that change during development.^4–7^ CSF has been shown to promote the function and survival of glia and neurons in vitro and may have a role in modulating neuronal activity.^7^

CSF flow has recently received renewed interest, in part due to the newly described pathways for fluid movement via the glymphatic and lymphatic pathways and their role in neurodegenerative diseases like Alzheimer’s Disease and multiple sclerosis.^8–10^ These routes for CSF movement have primarily been studied in adult and senescent animals, and the understanding of their role in the developing brain is still evolving. Historically implicated structures for CSF handling along the superior sagittal sinus, arachnoid villi and granulations, are not fully developed in humans until two years of age and are not present in a number of species including rodents.^11–13^ Glymphatic CSF handling has primarily been studied in adult animal models; most studies are focused on the superficial cortical surface adjacent to the middle cerebral artery (MCA) as it can be visualized with two-photon live imaging.^8, 14^ Finally, while sparse meningeal lymphatics have been observed in mice at birth around the foramen magnum and pterygopalatine artery^15^, most functional meningeal lymphatic connections take three to four weeks to fully develop postnatally.^15, 16^ CSF clearance via the Na-K-Cl (NKCC1) cotransporter during postnatal development was recently reported.^17^ However, this and other components of putative CSF movement, including lymphatic CSF outflow, glymphatic fluid and solute handling, perineural, perivenous and the spinal subarachnoid space (SAS) CSF handling have been studied largely in isolation from one another.^10, 18–20^ Few, if any, prior analyses of CSF dynamics have simultaneously been able to examine the cerebrum, cerebellum, and spinal cord due to imaging resolution, a limitation that hampers our ability to understand global CSF circulation throughout the central nervous system (CNS).

A significant challenge to the study of the pathologic CSF disturbances in pediatric hydrocephalus is the lack of understanding of how CSF (and its solutes) circulate within and out of the CNS during normal development. More specific is the concern that interruption of CSF generation and flow interrupts normal brain development. Posthemorrhagic hydrocephalus (PHH) after preterm intraventricular hemorrhage (IVH) is the most common cause of hydrocephalus in the United States and is closely linked to brain injury.^21^ The etiology for PHH is not known and studies of individual CSF pathways have not resulted in effective therapeutic targets or specific causes for the development of PHH.^22–26^ To pursue this question, we utilized a novel CSF imaging method, gold nanoparticle-enhanced X-ray microtomography (AuNP-XRM)^27–29^, for global, high-resolution 2-D and 3-D imaging of CSF pathways through the entire CNS with histologic confirmation.

Here, we present a comprehensive map of intraparenchymal CSF solute distribution using AuNP-XRM. Small CSF solutes travel via long projections from the basal cisterns to deep cholinergic neuronal cell populations and short projections to the cerebral and cerebellar cortices. Large CSF solutes target stem cell-rich populations via transependymal and transleptomeningeal routes. CSF solutes efflux via CNS borders including dura, arachnoid, choroid plexus, venous drainage, and cranial nerves. Finally, intraparenchymal CSF circulation is altered in PHH, with implications for mechanisms of brain injury and impaired development after IVH.

## Results

### Novel imaging of CSF circulation using AuNP-XRM

To image all pathways for CSF movement within the CNS, we utilized high resolution XRM with AuNPs as a CSF tracer.^27–29^ AuNP-XRM samples were prepared as shown in Figure 1 (see Methods). To generate the control and PHH conditions, a 20 µl solution of either artificial CSF (aCSF) or Hb was injected into the right lateral ventricle of post-natal day 4 (P4) rats. All animals in this study received either an aCSF or Hb injection at P4 unless otherwise indicated. At P7, 15-nm or 1.9-nm PEG-coated AuNPs constituted in aCSF were injected into the right lateral ventricle as a contrast agent and CSF tracer. The AuNPs have high X-ray attenuation and were verified to have a low negative zeta-potential and average hydrodynamic diameters of 35.91 and 3.79 nm for the 15-nm and 1.9-nm AuNPs, respectively (Figure 1d).^30, 31^ All injections were performed within two hours of noon to control for circadian effects on CSF circulation.^32, 33^ Rodents were returned to the cage to allow AuNPs to circulate within the CSF before perfusion and embedding for image acquisition. XRM images of a rodent without AuNP injection were used as a negative control. Image acquisition per animal ranged 12-18 hours in length. The use of this method resulted in ultra-high resolution (14.7 µm) images of the entire brain and spinal cord in which the CSF spaces, discrete nuclei (cell groupings), tracts, blood vessels, and nerves could be clearly visualized. To eliminate the possibility of imaging artifact from the perfusion process, separate samples were imaged after drop fixation with no difference in findings. To confirm AuNP distribution observed with AuNP-XRM, we harvested the brain and spinal cord post-XRM and processed the tissues for histology. Scanning electron microscopy (SEM) was also used to visualize AuNPs in the leptomeninges over the cerebral cortex.

**Figure 1.**
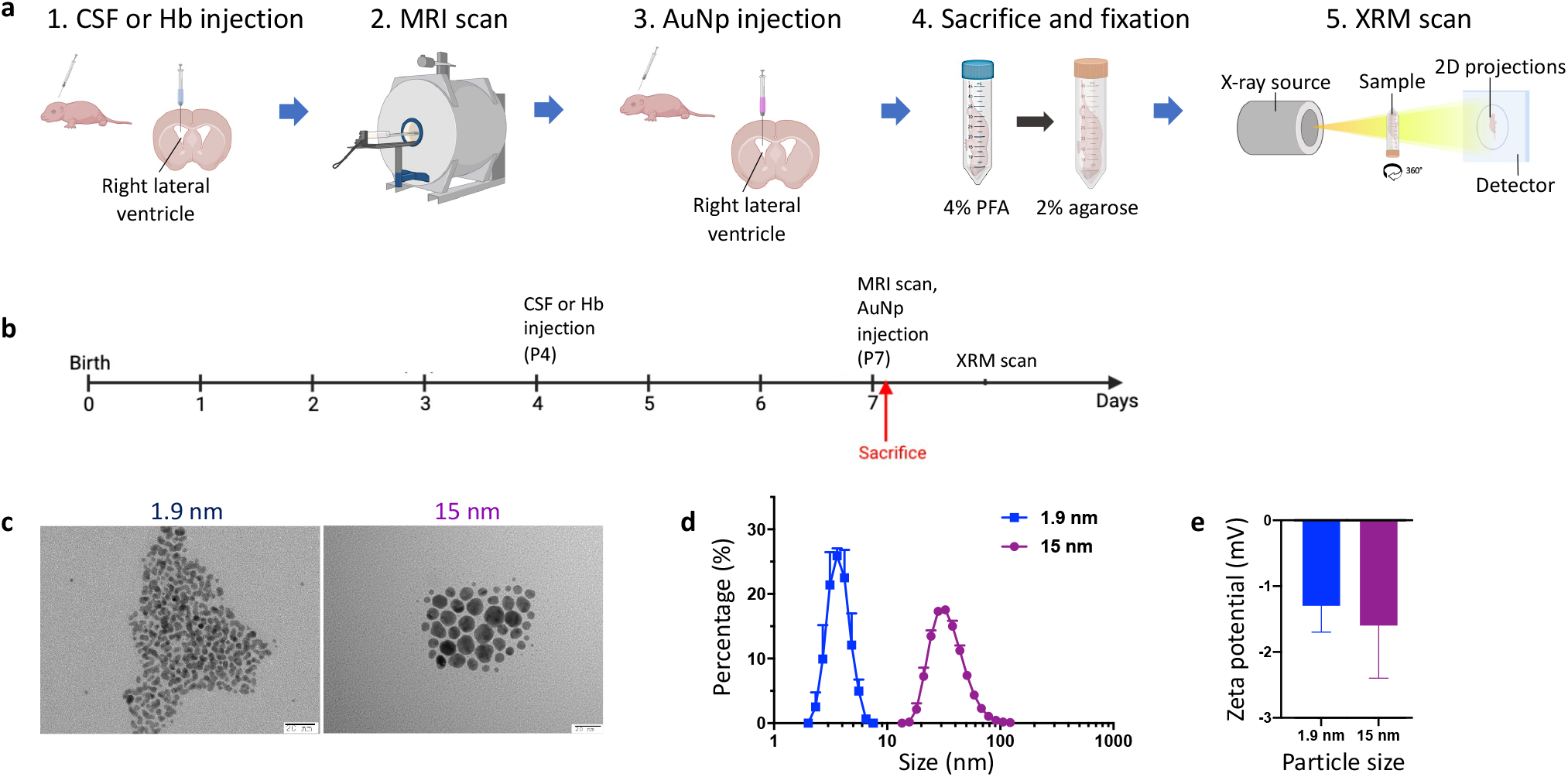
Gold nanoparticle enhanced X-ray microtomography (AuNP-XRM) **a-b**, Workflow (a) and timeline (b) for AuNP-XRM. P4 rodents were injected with artificial CSF (aCSF) or hemoglobin (Hb) into the right lateral ventricle to create the aCSF control and intraventricular hemorrhage (IVH) conditions. 72 hours later at P7, rodents underwent MRI scans to determine ventricle size and onset of post hemorrhagic hydrocephalus (PHH) following IVH. AuNPs were then injected into the lateral ventricles and allowed to circulate. Rodents were then perfused and fixed in agarose for XRM. **c-d,** Transmission electron microscopy (TEM) analysis (c) and size calculation (d) of 1.9-nm and 15-nm AuNPs. On average, the hydrodynamic diameter of 1.9-nm AuNPs was 3.79 nm while the hydrodynamic diameter of 15-nm AuNPs was 35.91 nm. **e,** 1.9 and 15-nm AuNPs have low zeta potentials (15-nm -1.6 ± 0.8 mV, 1.9-nm -1.3 ± 0.4 mV).

### Novel basal and dorsal CSF influx patterns

Unlike prior studies in adult mice in which CSF solutes were observed in dorsal structures^10, 34, 35^, we observed AuNPs primarily in deep and basal structures, with very little enhancement in and around the superior sagittal sinus (SSS) (Figure 2a, 2c). Following this observation, we hypothesized that CSF influx into the parenchyma occurs along more basal structures in neonatal rodents. Four hours after 15-nm AuNP injection, we observed a striking pattern of CSF projections formed by 15-nm AuNPs along the perivascular spaces of the vessels entering the brain around the interpeduncular fossa (Figure 3a, 3b). As early as 10 minutes post-1.9-nm AuNP injection, we observed 1.9-nm AuNPs along similar basal areas as we observed with 15-nm AuNPs (Figure 3d-f), forming intraparenchymal CSF projections. 1.9-nm AuNPs were also observed to travel from the SAS to the ventral horn of the spinal cord (Figure 3g) and dorsally over the convexities into layers III and V of the cortex (Figure 3g). These dorsal CSF projections were significantly shorter than those around the interpeduncular fossa (Figure 3h). In the spinal cord, projections to the ventral horn were significantly longer than projections to the dorsal horn (Figure 3i). The difference in dorsal convexity and spinal cord patterns observed between 1.9-nm AuNPs and 15-nm AuNPs indicate that these dorsal projections may be independent of penetrating vasculature. As expected, 15-nm AuNPs outlined perivascular spaces, and while we observed similar patterns of 1.9-nm AuNP pathways into the brain and spine, their morphology was similar to, but not clearly vascular for each individual projection when compared to a rat vascular atlas.^36^ In addition, there appeared to be more 1.9-nm AuNP CSF projections on the XRM than vessels identified with CD31 immunostaining, particularly around the interpeduncular fossa and dorsal convexities (Figure 3), suggesting that not all of the projections represent perivascular CSF movement.

**Figure 2.**
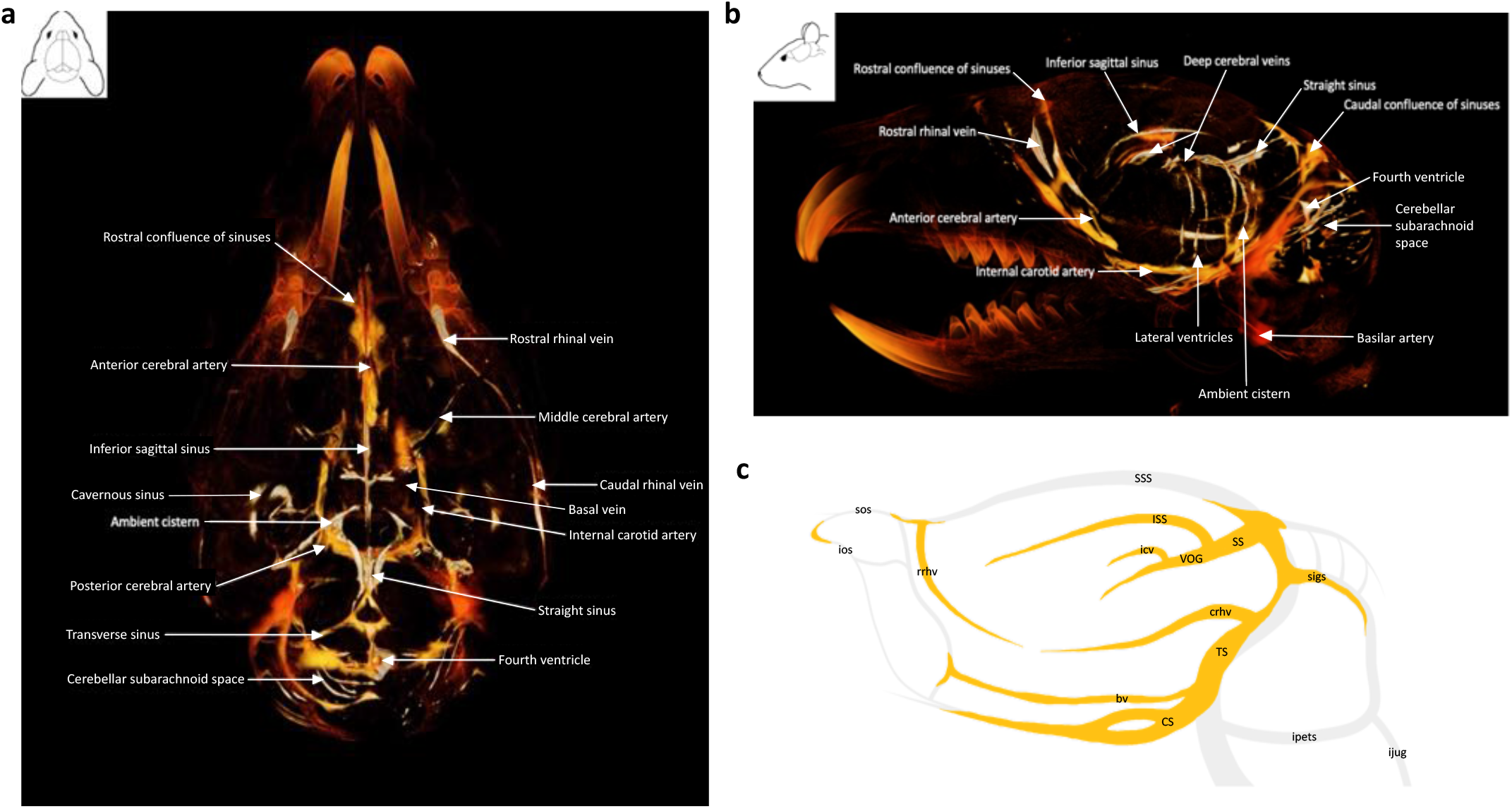
lntraventricularly injected CSF solutes primarily circulate through deep and basal structures in the rodent brain. **a-b**, Superior (a) and lateral (b) views of a 30 reconstruction of gold nanoparticle-enhanced X-ray microtomography (AuNP-XRM) showing distribution of AuNPs through the brain 4 hours after 15-nm AuNP injection into the right lateral ventricle in P7 aCSF control rodents. **c**, Schematic showing patterns of AuNP distribution observed in the venous system of the rat brain. Vasculature is indicated in gray, gold nanoparticle presence is indicated in gold. Abbreviations: sos, superior olfactory sinus; ios, inferior olfactory sinus; rrhv, rostral rhinal vein; SSS, superior sagittal sinus; ISS, inferior sagittal sinus; icv, internal cerebral veins; VOG, vein of Galen; SS, straight sinus; crhv, caudal rhinal vein; bv, basal vein; CS, cavernous sinus; TS, transverse sinus; sigs, sigmoid sinus; ipets, inferior petrosal sinus; ijug, internal jugular vein.

**Figure 3.**
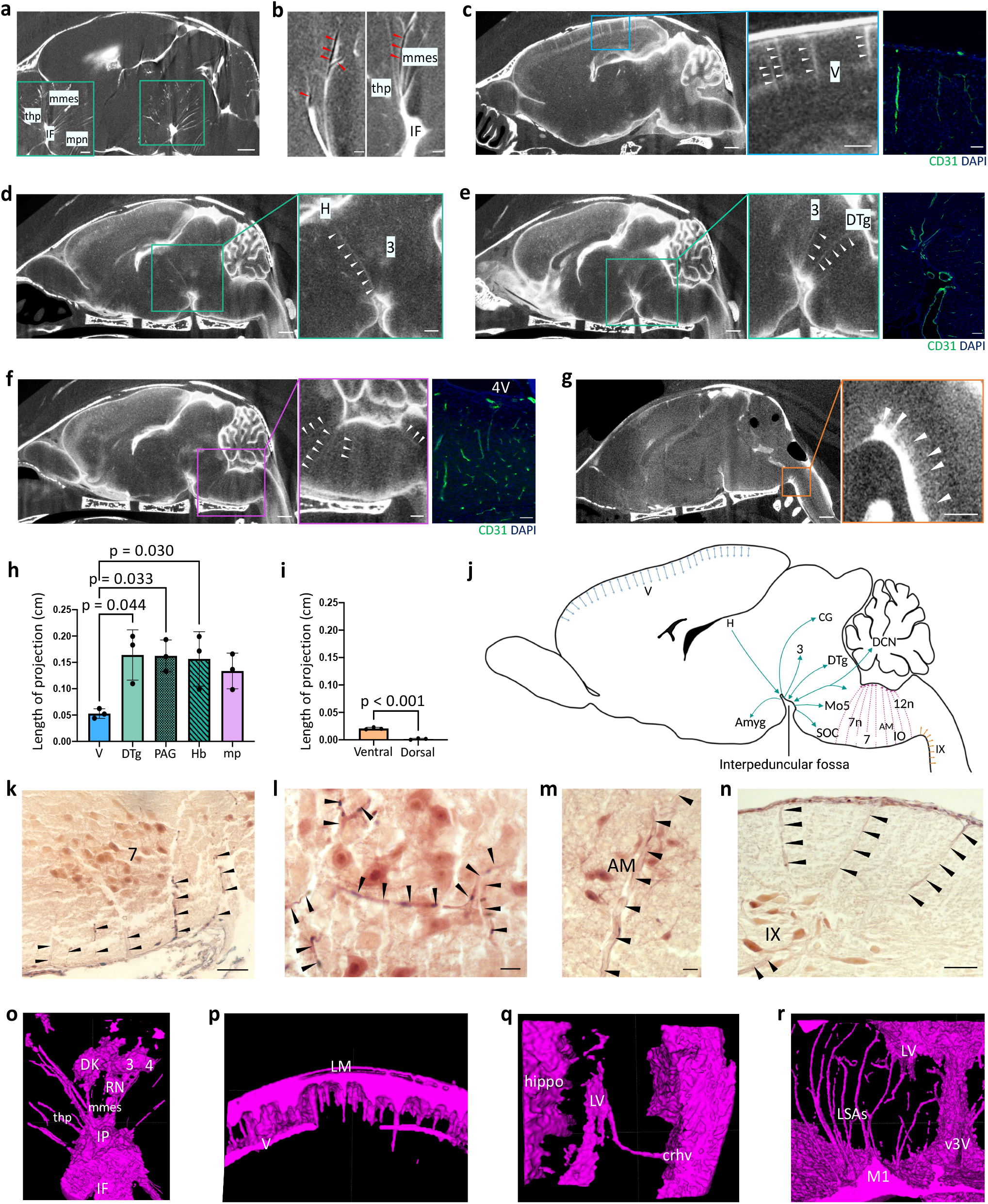
Basal and dorsal CSF projections target intraparenchymal nuclei throughout the forebrain, midbrain, and hindbrain. **a-b,** Sagittal X-ray microtomography (XRM) demonstrates 15-nm gold nanoparticle (AuNP) CSF influx along the perivascular spaces of the thalamoperforating (thp), median mesencephalic (mmes), and median pontine (mpn) arteries from the interpeduncular fossa (IF) 4 hours following injection at P7 in an aCSF control rodent. Lumen (dark, no gold) is indicated by red arrowheads. a scalebar = 1 mm, inset scalebar = 500 µm, b scalebar = 250 µm. **c-g,** CD31 immunostaining (right) and XRMs (left) demonstrating 1.9-nm AuNP CSF influx (white arrowheads) from the subarachnoid space (SAS) into layer V of the cerebral cortex (c), interpeduncular fossa into midbrain and pontine nuclei (d-e), pontine cistern into the brainstem (f), and spinal SAS into the ventral horn of the spinal cord (g). Sagittal XRM scalebars = 1 mm, inset scalebars = 100 µm, immunostaining scalebars = 50 µm. **h,** Quantification of 1.9-nm AuNP CSF influx depth into the brain parenchyma along basal and cortical projections. Data are mean ± s.e.m., n = 3 rodents, one-way ANOVA with post-hoc Tukey. **i,** Quantification of 1.9-nm AuNP CSF influx depth into the spinal cord parenchyma along ventral and dorsal projections. Data are mean ± s.e.m., n=3 rodents, unpaired two tailed t-test. **j,** Schematic representation of all observed AuNP CSF projections. **k-n,** Representative histology showing 1.9-nm AuNP CSF influx into the facial nucleus (k, i), nucleus ambiguus (m), and lamina IX (n). k scalebar = 100 µm, l scalebar = 25 µm, m scalebar = 25 µm, n scalebar = 100 µm. **o-r,** Semi-automatic segmentations showing 1.9 nm AuNP CSF influx at the interpeduncular fossa (o), cortex (p, q), and caudate putamen (r). o-q scalebar = 500 µm, r scalebar = 1 mm. All abbreviations indicated in Supplementary Table 3.

### Small CSF solutes target specific neuronal populations in the brain and spinal cord

CSF distribution within the brain has previously been described as a global process whereby CSF and the brain itself are considered single entities.^37–39^ As the brain has a diverse range of cell types and networks that coordinate regional and global network functions^40, 41^, we hypothesized that in addition to a diffuse, global interstitial glymphatic CSF circulation, there exists a targeted CSF circulation within the brain parenchyma coordinated by the CSF projections from the interpeduncular fossa and SAS. At the termination of these CSF projections into the brain and spinal cord, we observed focal, intense regions of 1.9-nm AuNPs which corresponded to specific cell groupings in the forebrain, midbrain, brainstem and cerebellum.^42, 43^ For example, interpeduncular projections terminated in the habenula (H), oculomotor nucleus (3 CN nucleus) and dorsal tegmentum (DTg); cortical SAS projections terminated in layers III and V of the cortex; and spinal SAS projections terminated in lamina IX (IX) in the ventral horn (Figure 3a-g, o-p). We quantified the signal intensity in each anatomic area observed to have 1.9-nm AuNPs against the signal of the surrounding background across 3 rodents and identified 24 nuclei and cell groupings with a significant increase in 1.9-nm AuNPs compared to the background (Figure 4a-d, Extended Data Figure 1). There were also three nuclei (red nucleus, nucleus of Darkschewitsch, trochlear nucleus) with increased 1.9-nm AuNPs in one rodent, and one nucleus (nucleus sagulum) with increased 1.9-nm AuNPs in two rodents. In addition, we observed widespread distribution of 1.9-nm AuNPs in the molecular layer of the cerebellum (Extended Data Figure 2). In the spinal cord, 1.9-nm AuNPs were observed in lamina IX of all rodents, suggesting preferential CSF flow into the ventral horn (Figure 4, Extended Data Figure 1w). In addition to the targeted distribution to these 24 nuclei, there was a diffuse, global distribution of 1.9-nm AuNPs to the parenchyma at 10 minutes, compared to our negative control XRM without background signal. The corpus callosum (anterior forceps, body, genu, and splenium) was the only brain region identified that did not have 1.9-nm AuNP distribution or uptake, which highlights the neuronal and grey matter preferential handling of CSF. We show early, specific 1.9-nm AuNP distribution to neuronally rich regions of the brain, spinal cord, and specifically the cerebellum with implications for CSF uptake and handling in these regions which may differ from other brain regions. To our knowledge, this is the first detailed description of CNS wide CSF interaction with specific cell populations within the brain, including significant cerebellar CSF distribution.

**Figure 4.**
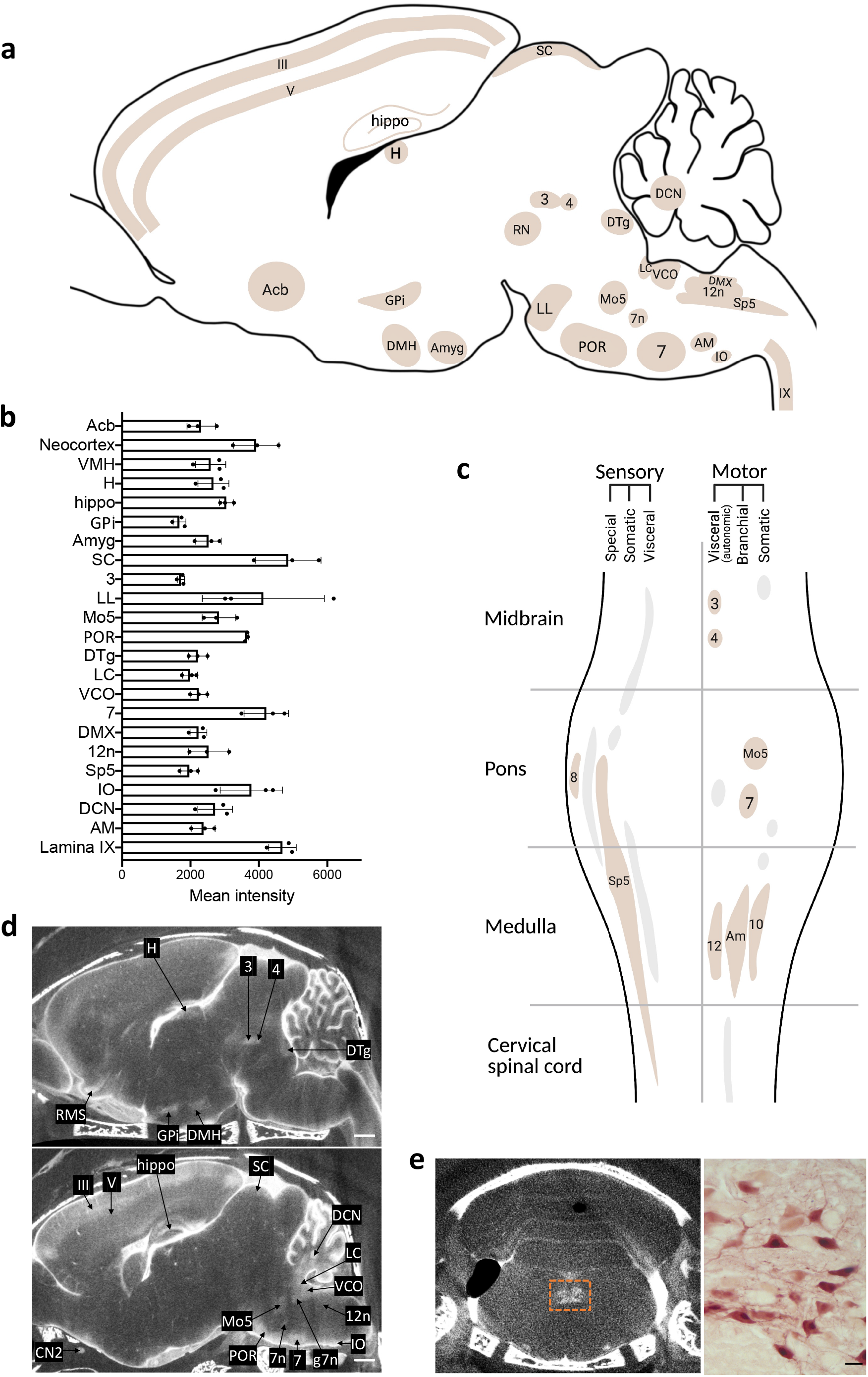
Neuronal and stem cell distribution of small CSF solutes in the brain and spinal cord parenchyma. **a,** Schematic representation of nuclei and other cell groupings in which 1.9-nm gold nanoparticles (AuNPs) were observed 10 minutes after intraventricular injection in P7 control rodents. Areas identified in all rodents include the nucleus accumbens (Acb), layers III and V of the neocortex (III, V), dorsal medial hypothalamus (DMH), habenula (H), hippocampus (hippo), globus pallidus internus (GPi), amygdala (Amyg), interpeduncular nucleus (IP), superior colliculus (SC), oculomotor nucleus (3), ventral lateral lemniscus (VLL), motor trigeminal nucleus (Mo5), periolivary region (POR), dorsal tegmental nucleus (DTg), locus coeruleus (LC), ventral cochlear nucleus (VCO), facial nucleus (7), dorsal nucleus of the vagus nerve (DMX), hypoglossal nucleus (12n), spinal trigeminal nucleus (Sp5), inferior olivary nucleus (IO), deep cerebellar nuclei (DCN), nucleus ambiguus (AM), and lamina IX of the spinal cord (IX). 1.9-nm AuNPs were identified in the nucleus of Darkschewitsch (DK), red nucleus (RN), trochlear nucleus (4) in one rodent and the nucleus sagulum (SAG) in two rodents. **b,** Quantification of 1.9-nm AuNP mean intensity in cell groupings. **c,** Schematic representation of 1.9-nm AuNP distribution in cranial nerve nuclei. Nuclei with 1.9-nm AuNP uptake (brown) are primarily motor. **d,** Representative X-ray microtomography (XRM) images showing 1.9-nm AuNP distribution. scalebars = 1 mm. **e,** Representative coronal XRM (left) and histology (right) showing 1.9-nm AuNP distribution in the 12n (orange box, inset). XRM scalebar = 1 mm, histology scalebar = 25 µm. **a-e** Additional analyses shown in Extended Data Figure 1. All data representative from three rodents.

We used four methods to validate our observations. First, 1.9-nm AuNP-XRM images were registered to the Duke Wistar rat atlas^44^ using 4dfp tools (http://4dfp.readthedocs.io); skulls were stripped and regions of intraparenchymal CSF were semi-automatically segmented to reveal the same patterns of CSF distribution, suggesting our observations were not the result of computer artifact. Second, to confirm cell type and location within the anatomic regions identified on 1.9-nm AuNP-XRM, the brain and spinal cord were histologically sectioned without any additional markers or counterstains. 1.9-nm AuNPs, which appear brown under light microscopy, were observed within neurons in the cell groupings that corresponded to the same anatomic locations seen on AuNP-XRM (Extended Data Figure 1). Cresyl violet counterstaining confirmed that a subpopulation of neurons (i.e., not all neurons) within each anatomic cell grouping contained gold ranging from 12-67% of all neurons, suggesting a unique CSF uptake function for neurons with vs. without gold (Extended Data Figure 3), and confirming that the that CSF uptake within these cell groupings was not uniform across all neurons.

Third, to confirm that the observed CSF distribution patterns were independent of the physical and chemical characteristics of the 1.9-nm AuNPs and to confirm similar distributions with in vivo imaging, we acquired in vivo intraventricular gadolinium-enhanced MRIs (5 mmol/mL Dotarem, Guerbet, Princeton, NJ, USA) (Extended Data Figure 4) and observed high concentrations of gadolinium 30 minutes post gadolinium injection in cell groupings similar to those observed with 1.9-nm AuNPs (Extended Data Figure 4). These similar patterns suggest that the 1.9-nm AuNP distribution is not an artifact of the ex vivo nature of XRM or specific to AuNPs, but rather generalizable to small CSF solutes.

Fourth, to evaluate cross-species preference of small CSF solutes for the 24 identified neuronal cell populations, we performed intraventricular injection of Red-Dextran (70kDa, R9379, Millipore Sigma, Burlington, MA, USA) into the ventricles of 4 days post-fertilization (DPF) zebrafish before imaging at 5 DPF. Similar to the CSF distribution patterns of 1.9-nm AuNPs and gadolinium, Red-Dextran was observed in neuron-rich populations throughout the zebrafish brain (Extended Data Figure 5).

**Figure 5.**
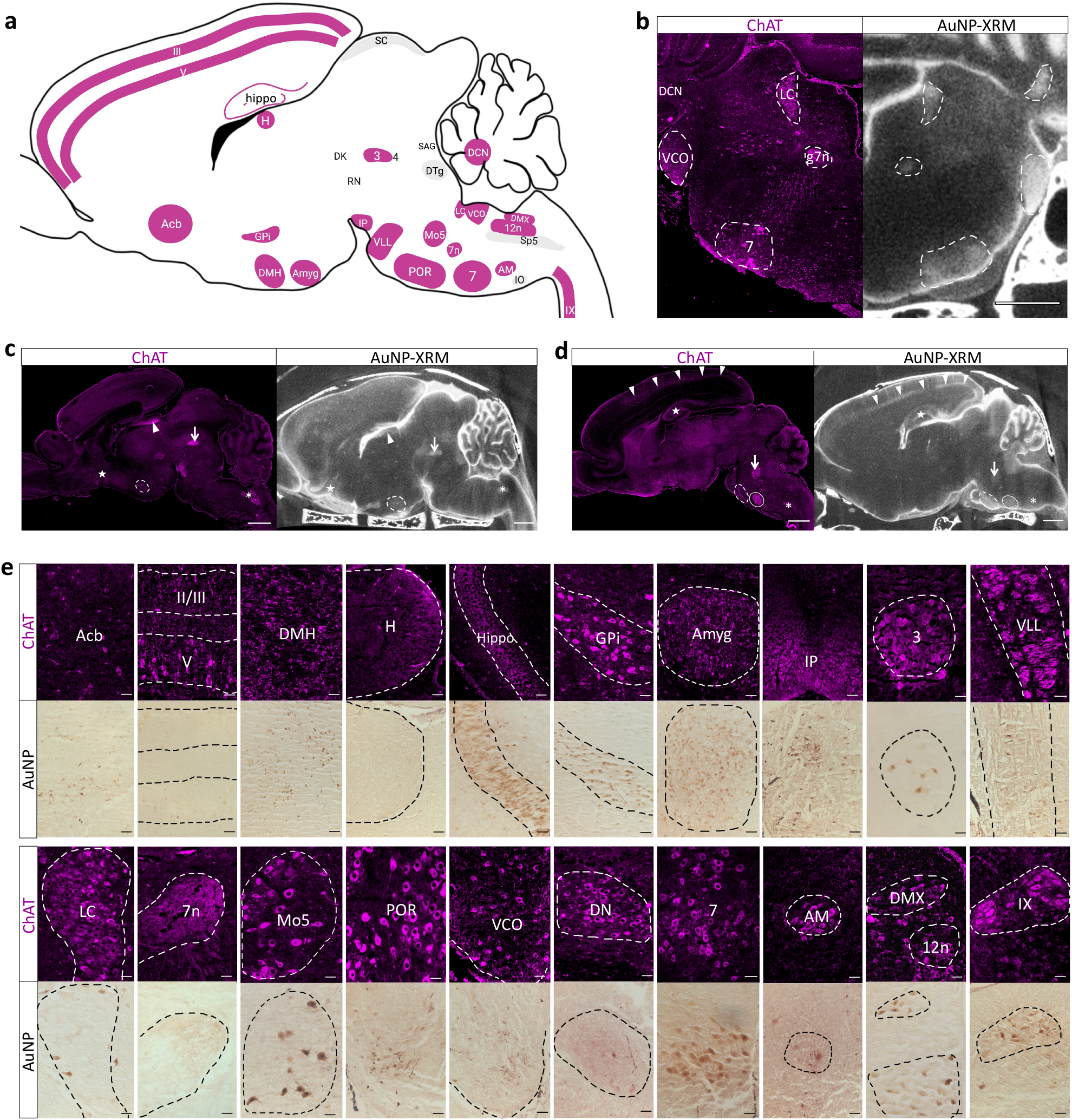
CSF distribution mirrors choline acetyltransferase (ChAT) expression in the rodent brain. **a,** Schematic representation ChAT-expressing brain nuclei and cell groupings in which 1.9-nm gold nanoparticles (AuNPs) were observed 10 minutes after intraventricular injection at P7 in an aCSF control rodent. Four nuclei, the superior colliculus (SC), dorsal tegmentum (DTg), spinal trigeminal nucleus (Sp5), inferior olive (1O), with AuNP distribution were not observed to have ChAT expression. Abbreviations: nucleus accumbens (Acb), neocortex (11/111, V), dorsal medial hypothalamus (DMH), habenula (H), hippocampus (hippo), globus pallidus internus (GPi), amygdala (Amyg), interpenduncular nucleus (1P), oculomotor nucleus (3), ventral lateral lemniscus (VLL); locus coeruleus (LC), genu of the facial nerve (7n), motor trigeminal nucleus (Mo5), periolivary region (POR), ventral cochlear nucleus (VCO), dentate nucleus (DN), facial nerve nucleus (7), nucleus ambiguus (AM), dorsal motor nucleus of the vagus nerve (DMX), hypoglossal nucleus (12n), lamina IX of the spinal cord (IX). **b-e**, Representative coronal (b), sagittal (c-d) and higher magnification (e) immunofluorescence sections showing ChAT expression (left) and AuNP distribution on XRM (right). C, star, Acb; dotted circle, DMH; arrowhead, H; arrow, 3; asterisk, AM. d arrowheads, neocortex; star, hippo; arrow, Mo5; dotted circle, POR; circle, 7. b-d scalebars = 1 mm, e scalebars = 50 µm. ChAT expression in the nucleus of Darkschewitsch (DK), red nucleus (RN), trochlear nucleus (4) in one rodent and the nucleus sagulum (SAG) was not examined because 1.9-nm AuNP distribution was not observed in those regions in all animals. Data is representative from 3 rodents.

Finally, through direct review of figures in published reports of either CSF tracers or solutes administered to the CSF for other purposes, we confirm that the patterns and cell grouping identified here with high resolution 1.9-nm AuNP-XRM are consistent with existing studies, although these prior studies had not specifically noted these patterns.^45, 46^ Furthermore, previous studies using intraventricular injection of radioactive transferrin in rodents demonstrated similar patterns of transferrin distribution to the 1.9-nm AuNP distribution^47^; another study in which intrathecal injection of adeno-associated virus capsid serotype 9 (AAV9) for transgene delivery transduced neurons in some of the same nuclei in which AuNPs were observed.^48^ A third study showed intra cisterna magna [14C]inulin also had similar distribution patterns.^49^ There was also evidence for small fluorescent tracer distribution along the same interpeduncular fossa projections and brainstem nuclei identified with 1.9 nm AuNPs in a previous study. The observed 1.9-nm AuNP distribution patterns, combined with evidence from in vivo gadolinium MRI and zebrafish Red-Dextran studies, as well transferrin, AAV9, [14C]inulin, and 3kDa FITC-dextran distribution patterns suggest that there is differential intraparenchymal CSF distribution to specific cell groupings in the CNS.

While CSF handling along the MCA and SSS has received intense focus, we did not observe concentrated regions of AuNP delivery at the terminations of projections from the MCA (specifically the lenticulostriate arteries into the caudate putamen) (Figure 3r). We also did not observe a clear route from MCA cortical projections to the 24 neuronal cell populations, particularly those in the brainstem (Figure 3r). As small CSF solutes (1.9-nm AuNP, gadolinium, Red-Dextran) were present in high concentrations in the parenchymal neuronal cell populations, but not the adjacent brain regions, we hypothesized that the CSF takes a direct path to the 24 neuronal cell populations via specific CSF projections from the SAS including perivascular flow patterns. The localization is in contrast to either selective uptake into these cell populations from a uniform, global CSF influx or a gradient distribution. These findings suggest that these projections play a role in targeted delivery of CSF to specific intraparenchymal regions that may be independent of previously described glymphatic CSF transport.

### CSF distribution mirrors choline acetyltransferase expression in the brain

Given the diverse anatomic locations of observed targeted CSF delivery of small CSF solutes, we hypothesized that this anatomic and cell type-specific pattern of CSF distribution was mediated through the expression of genes, neurotransmission, or function. Notably, seven of nine cranial nerve nuclei with concentrated small CSF solutes were motor-derived cranial nerve nuclei (Figure 4c), and the other 14 cell groupings had roles in motor, learning, and memory functions, which suggests a common developmental origin for the areas of the brain preferentially receiving CSF projections. We systematically reviewed the Gene Expression Nervous System Atlas (GENSAT)^50^ and the Allen Mouse Brain Atlas Anatomic Gene Expression Atlas (AGEA)^51^ of the developing brain to identify similar gene expression patterns to that of the observed CSF distribution patterns (Figure 4) and found strikingly similar patterns to that of choline acetyltransferase (ChAT) (Figure 5, Supplementary Tables 1, 2). Microscopy imaging co-stain of tissue sections for ChAT and AuNPs was not possible as AuNPs blocked transmitted light.

Therefore, histologic sections from a separate cohort of aCSF control rodents without AuNP injection at P7 were stained for ChAT. ChAT immunofluorescence showed highly similar staining patterns within 20 of 24 of the same cell groupings as the 1.9-nm AuNPs (n=3 rodents) (Figure 5). Cross reference with an atlas (GENSAT) and previously published data supported parallel ChAT expression to CSF circulation patterns.^50^

### Transependymal and transleptomeningeal circulation of large CSF solutes

The size of the space between overlapping astrocyte end feet allowing passage of CSF and its solutes from the perivascular spaces into the parenchyma is accepted to be approximately 20 nm.^8, 52^ Accordingly, we reasoned that large CSF solutes (15-nm AuNP core, with 35.91 nm hydrodynamic diameter) could enter the perivascular space (Figure 2, 3a, 3b), but not subsequently efflux into the ISF to distribute through the parenchyma. This finding would therefore distinguish between perivascular and non-perivascular influx dependent intraparenchymal CSF circulation patterns.

Ten minutes after intraventricular injection of 15-nm AuNPs in wild type animals (no aCSF injection at P4), we observed AuNPs unilaterally in the right olfactory bulb through the rostral migratory stream (RMS), subventricular zone (SVZ) and corpus callosum (Figure 6a, 6b). There was more widespread distribution at 4 hours (Figure 6b), with 15-nm AuNPs observed bilaterally in the RMS, SVZ, corpus callosum, as well as the brain parenchyma medial to the lateral ventricles including the hippocampus (Figure 6b). In contrast to the distribution of small CSF solutes through cortical projections, there was not significant 15 nm AuNP intraparenchymal influx from the surface (Figure 6c). Instead, 15-nm AuNPs extended to the parenchyma in a gradient distribution from the ventricular ependyma in a region-specific manner, with the highest concentration near the medial and superior wall of the lateral ventricles with decreasing concentration superiorly into the corpus callosum and medially into the lateral septal nucleus (Figure 6d-g). A similar gradient was observed from the superior wall of the lateral ventricle into the corpus callosum and the roof of the third ventricle into the hippocampus, suggesting these large particles were transported via non-perivascular transependymal routes into the brain parenchyma (Figure 6d-g, 5l, 5k). Notably, the transependymal movement of large CSF solute was not uniform across the ventricle walls with preferential flow across the ependyma of the medial and superior walls of the lateral ventricle over the lateral wall (Figure 6d-g, 6h).

**Figure 6.**
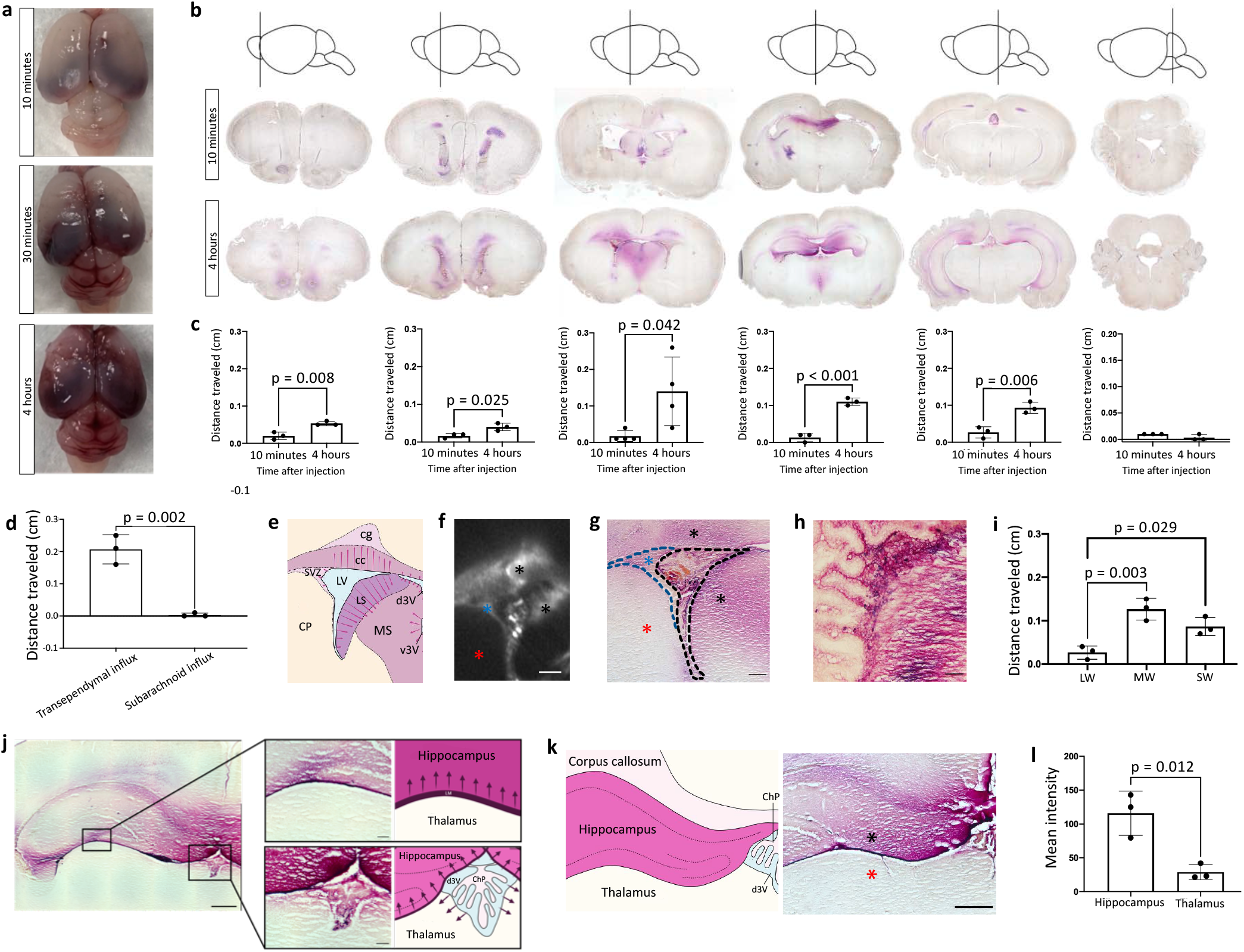
Region-specific perivascular-independent transependymal circulation of large CSF solutes. **a-b,** Representative gross (a) and histology (b) images of 15-nm gold nanoparticles (AuNP) (purple) distribution 10 minutes, 30 minutes, and 4 hours after intraventricular injection in P7 rodents. **c,** Quantification of distance traveled by 15-nm AuNPs from the center of the olfactory bulb (OB), rostral migratory stream (RMS) and the ependymal surface of the medial wall of the lateral ventricle, ventral third ventricle, dorsal third ventricle, and floor of the fourth ventricle at 10 minutes and 4 hours post-intraventricular injection. Data are mean ± s.e.m., n = 3 rodents, unpaired two-tailed t-test. **d,** Quantification of distance traveled by intraventricularly injected 15-nm AuNPs into the brain parenchyma transpendymally compared to influx from the subarachnoid space. Data are mean ± s.e.m., n = 3 rodents, unpaired two-tailed t-test. **e-g,** Schematic (e), representative X-ray microtomography (XRM) (f), and histology (g) demonstrating transependymal 15-nm AuNP circulation through the medial and superior walls (black asterisks) and the subventricular zone (SVZ) (blue asterisk) of the right lateral ventricle. Little 15-nm AuNP distribution was observed in the lateral wall (red asterisk). Scalebars = 250 µm. Abbreviations: SVZ, subventricular zone; CP, caudate putamen; cg, cingulum; cc, corpus callosum; LS, lateral septal nucleus; MS, medial septal nucleus; d3V, dorsal third ventricle; v3V, ventral third ventricle; LV, lateral ventricle. **h,** Higher magnification image of right lateral ventricle choroid plexus and medial wall. Scalebar = 50 µm. **i,** Quantification of distance traveled by 15-nm AuNPs from the medial wall (MW), superior wall (SW) and lateral wall (LW). Data are mean ± s.e.m., n = 3 rodents, one-way ANOVA with post-hoc Tukey. **j,** Representative histology and schematics demonstrating 15-nm AuNP circulation into the. scale bar = 500 µm, inset scalebar = 100 µm) **k,** Schematic and histology demonstrating 15-nm AuNP transependymal circulation through the dorsal the third ventricle into the hippocampus (black asterisk). Very little 15-nm AuNP circulation was observed in the thalamus (red asterisk). scalebar = 1000 µm. **l,** Quantification of intensity of 15-nm AuNPs in the hippocampus and thalamus. Data are mean ± s.e.m., n = 3 rodents, unpaired two-tailed t-test) **d-l,** all observations and quantifications were made 4 hours post-15-nm AuNP injection into the right lateral ventricle. All data are representative from three rodents.

Most notably, 15-nm AuNPs entered the hippocampus through two routes: transependymal across the roof of the third ventricle and from the leptomeninges of the quadrigeminal cistern (Figure 6l). Strikingly, 15-nm AuNPs originating from the leptomeninges of the perimesencephalic cisterns were only observed to extend superiorly into the hippocampus without any movement inferiorly into the thalamus (Figure 6j, 6k). This transependymal and transleptomeningeal-hippocampal movement of 15-nm AuNPs may have implications for the clearance of large CSF solutes, including large protein aggregates in neurodegenerative diseases such as Alzheimer’s disease.^53–55^

A direct histologic comparison 10 minutes after intraventricular injection of small and large CSF solutes revealed distinct differences in solute handling within a given area of the brain/spine. As expected, 1.9-nm AuNPs covered significantly more total cross-sectional brain area than 15-nm AuNPs (Figure 7a, 7b). In the hippocampus, midbrain, brainstem, and spinal cord, very little 15-nm AuNP signal was observed in the parenchyma 10 minutes after injection, compared to diffuse 1.9-nm AuNP signal throughout the parenchyma with additional concentrated AuNPs in the previously mentioned nuclei (Figure 7d-f). Within the choroid plexus (CP), 15-nm AuNPs were observed on the apical surface of the lateral ventricle CP, while 1.9-nm AuNPs were observed on the apical and basal surfaces of the lateral ventricle CP (Figure 6g).

**Figure 7.**
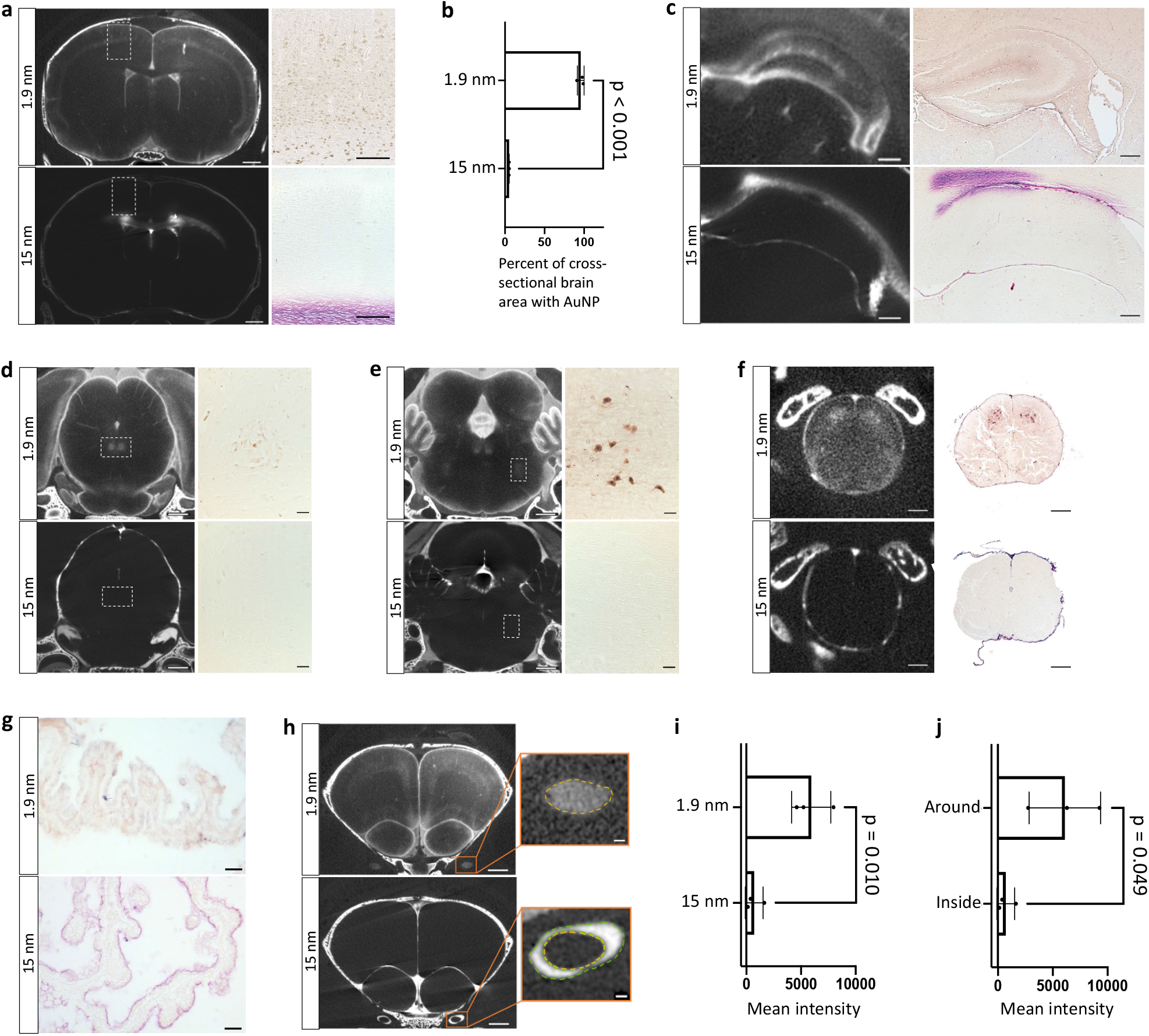
Size-dependent CSF solute distribution in the parenchyma, choroid plexus, and optic nerve. **a,** Representative X-ray microtomography (XRM) images of 1.9-nm (top) and 15-nm (bottom) gold nanoparticle (AuNP) distribution 10 minutes after intraventricular injection in P7 aCSF control rodents. Histology showing AuNP distribution in neurons of the neocortex (1.9-nm) and supraventricular white matter (15-nm). scalebar = 1 mm, inset scalebar = 100 µm. **b,** Quantification of percentage of the cross-sectional brain area with 1.9-nm and 15-nm AuNP circulation. Data are mean ± s.e.m., n = 3 rodents, unpaired two-tailed t-test. **c-f,** Representative XRM images and histology of 1.9-nm (top) and 15-nm (bottom) AuNP distribution in the hippocampus (c), midbrain and oculomotor nucleus (d), pons and motor nucleus of the trigeminal nerve (e), and spinal cord (f). c scalebars = 250 mm, d-e scalebar = 1 mm, d-e inset scalebar = 50 µm, f scalebars = 250 µm. **g,** Representative histology of 1.9-nm (top) and 15-nm (bottom) AuNP distribution in the right lateral ventricle choroid plexus. scalebar = 25 µm. **h-j,** Representative XRM images of 1.9-nm (top) and 15-nm (bottom) AuNP distribution in the optic nerve (h) and quantification of the mean intensity of 1.9-nm and 15-nm gold nanoparticles within the optic nerve (i). There was also a significant difference in the mean intensity of 15-nm gold nanoparticles inside and around the optic nerve (j). g scalebar = 1 mm, g inset scalebar = 100 µm. All observations and quantifications were made 10 minutes post-1.9-nm and 15-nm AuNP injection into the right lateral ventricle of P7 aCSF control rodents. i-j, Data are mean ± s.e.m., n = 3 rodents, unpaired two-tailed t-test)

### Meningeal handling of large CSF solutes in the brain and spinal cord

Next, we examined differential handling of CSF by the cranial leptomeninges (pia-arachnoid) and dura. We observed limited 15-nm AuNP circulation within the dura on a meningeal wholemount, however on a cross section of the brain we observed widespread 15-nm AuNP circulation through the leptomeninges of the perimesencephalic cisterns (Extended Data Figure 6a-k). On the cross sections, 15-nm

AuNPs were also observed in the pia traversing the CSF spaces intracranially, including the tela choroidea in the roof of the third ventricle (3V) and the lateral ventricle, 3V, and fourth ventricle CP (Extended Data Figure 6l-q). These patterns of 15-nm AuNP distribution through the tela choroidea were highly similar to a previously described pattern of superficial siderosis and has implications for direct communication between the ventricles outside the ependymal-lined CSF cavities.^56^ 15-nm AuNPs were also observed in the leptomeninges in the folia of the cerebellum (Extended Data Figure 6r). Distribution of 1.9-nm AuNPs was more diffuse through the dura compared to the 15-nm AuNPs, however the cross sections showed similar patterns of 1.9-nm and 15-nm AuNP distribution through the intracranial leptomeninges (Extended Data Figure 6).

We also evaluated large CSF solute distribution through the spine meninges. We observed little evidence for 15-nm AuNPs in the dura, except along the nerve roots exiting the spinal cord (Extended Data Figure 7a). In contrast, we observed widespread 15-nm AuNP circulation through the leptomeninges that outlined the structure of the median fissures and nerve roots (Extended Data Figure 7b, 7c). We also observed different patterns of 15-nm AuNP distribution along the ventral and dorsal nerve roots. (Extended Data Figure 7b, 7c)

### Differential distribution of large and small CSF solutes in and around the optic nerve.

Similar to prior studies, we observed CSF in and around cranial nerves. It has previously been suggested that cranial nerves play a role in CSF outflow to the lymphatic system^15, 18, 57^, where a recent study of human lymphatic CNS elements found LYVE-1 staining in the endoneurium of the trigeminal nerve and the perineurium of the glossopharyngeal, vagus, and accessory cranial nerves.^57^ By comparing small and large CSF solute distribution, we find large CSF solute circulation around the optic nerve as it emerged from the chiasm, into the eye (Figure 7h), compared to small CSF solutes, which were observed within the optic nerve itself (Figure 7h, 7j).

### Targeted CSF distribution is altered in neonatal posthemorrhagic hydrocephalus

Our findings of targeted CSF delivery to cholinergic and stem-cell rich cell populations suggests a functional role of the basal CSF projections in brain development. Therefore, we hypothesized that altering CSF flow by inducing hydrocephalus would lead to specific changes in the targeted CSF pathways identified with AuNP-XRM, but not the diffuse interstitial distribution. To test this hypothesis, we induced IVH-PHH with hemoglobin injection into the right lateral ventricle of P4 rodents where aCSF-injected rodents served as controls.^59^ Ventricle volume was quantified using MR images acquired 72 hours post-injection with a significant enlargement in ventricle size in PHH (5.03±1.38 mm^3^) compared to aCSF controls (0.38±0.14 mm^3^) (Figure 8a, 8b).

**Figure 8.**
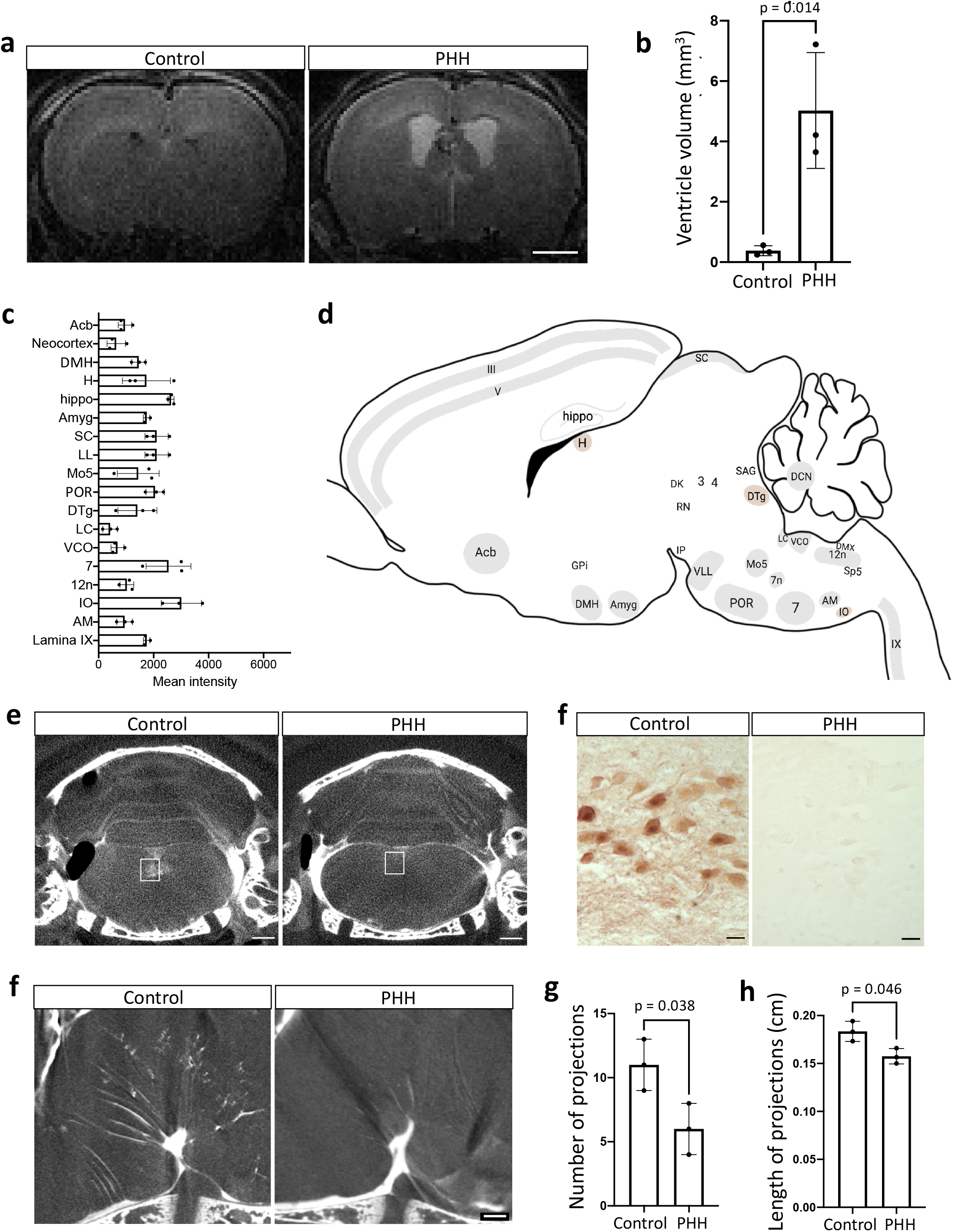
Targeted CSF distribution is altered in post hemorrhagic hydrocephalus {PHH). **a,** MRI demonstrating ventriculomegaly 72 hours after hemoglobin injection into the lateral ventricle of P4 rodents to create the PHH condition. **b,** Lateral ventricle volume was significantly increased in the PHH cohort compared to the aCSF controls. Data are mean ± s.e.m., n = 3 rodents, unpaired two-tailed t test. **c,** Quantification of 1.9-nm gold nanoparticle (AuNP) mean intensity in 18 nuclei and cell groupings in PHH rodents 10 minutes after intraventricular injection at P7. **d,** Schematic representation of brain nuclei and other cell groupings with significantly decreased 1.9-nm AuNP intensity (compared to aCSF control rodents) (gray). Cell groupings in which no significant changes in AuNP intensity were observed are shown in brown; those that were not able to be accurately identified or were not identified in all rodents are in white. **e-f,** Representative coronal X-ray microtomography (XRM) (e) and histology (f) showing AuNP distribution in the hypoglossal nucleus (12n) (white box) in aCSF control and PHH rodents. e scalebar = 1 mm, f scalebar = 25 µm. **g-h,** Representative XRM and quantifications demonstrating changes in the density and depth of influx of AuNPs along the basal projections. This corresponds to the decrease in AuNP distribution in the intraparenchymal nuclei. g scalebar = 100 µm. Data are mean ± s.e.m., n = 3 rodents, unpaired two-tailed t-test. **a-h** Coronal XRM, histology and quantifications of the other 17 regions were also performed (Extended Data Figure 8). All data representative from three rodents. All abbreviations indicated in Supplementary Table 4.

We found that while the diffuse interstitial 1.9-nm AuNP CSF distribution was present in both PHH and aCSF control animals, PHH was associated with significantly decreased 1.9-nm AuNP circulation in 16 of 24 cell groupings we originally identified to have targeted CSF distribution and concentrated 1.9 nm AuNPs. (Figure 8c, Extended Data Figure 8a-c, e-j, l-o, q, r). There were also non-significant differences in the 1.9-nm AuNP distribution in 3 of 24 nuclei (Extended Data Figure 8d, 8k, 8p). 1.9-nm AuNP distribution between aCSF control and PHH conditions in the remaining 5 nuclei (globus pallidus internus, interpeduncular nucleus, oculomotor nucleus, dorsal motor nucleus of the vagus nerve, spinal trigeminal nucleus) were not compared because the nuclei were not easily identifiable in the PHH group. Furthermore, because we observed perivascular CSF projections along branches of the PICA and BA reaching these nuclei, we evaluated the number and length of these CSF basal projections in the PHH condition compared to the normal brain. We observed significantly fewer projections from the interpeduncular fossa (Figure 8g, 8h), and that the remaining projections were significantly shorter in PHH compared to controls (Figure 8g, 8i).

The finding of IVH-PHH altering intraparenchymal CSF distribution to neuronal cell populations important in learning, memory, motor functions, has significant implications for CSF-mediated direct brain injury through transport of molecules/signals to neuronal populations involved in critical plastic functions. The connection between CSF movement (and potentially blood breakdown products like hemoglobin) into stem cell populations and the developmental association between the ventricles and their anatomical proximity to stem cell-rich brain regions like the hippocampus and SVZ merits further study.

In contrast to the 1.9-nm AuNP distribution in PHH, 15-nm AuNP distribution was increased in the olfactory bulb and corpus callosum, but there were no significant changes in 15-nm AuNP distribution in the cerebellum, SVZ and hippocampus (Extended Data Figure 9). These changes may be due to increased transependymal CSF flow across the lateral ventricle walls due to PHH-induced damage. Further longitudinal studies should be conducted to examine the long-term consequences of increased CSF solute circulation into these regions following PHH.

## Discussion

In this study, we used AuNP-XRM as a novel high resolution XRM-based CSF imaging method using 1.9-nm and 15-nm AuNPs to image CSF pathways through the brain and spinal cord in postnatal rodents. Without requiring extensive tissue processing, AuNP-XRM identified multiple novel CSF projections into the dorsal and basal aspects of the brain, the latter of which is an area that has not been extensively explored due to imaging constraints. This strategy also exploits the high resolution of X-rays and high X-ray attenuation of AuNPs to identify specific, novel cell populations to which CSF is targeted via the projections, including basal CSF influx to cholinergic cell populations, as well as other routes such as diffuse CSF influx to the cerebellar molecular layer and perivascular-independent transependymal circulation to stem cell rich populations such as the hippocampus.

CSF distribution has previously been thought to correlate with aquaporin-4 (AQ-4) expression patterns.^8, 60^ In this study, we propose a biological, neurotransmitter-driven intraparenchymal CSF-ISF circulation. Specifically, ChAT is a catalyst in the biosynthesis of acetylcholine (ACh)^61^, a widely distributed neurotransmitter involved in normal brain function and in the pathophysiology of inflammation and Alzheimer’s Disease.^62–69^ Moreover, CSF pathophysiology has been implicated in Alzheimer’s Disease and aging more broadly, where a reduction of CSF influx has been associated with Alzheimer’s Disease and enhanced function of putative routes of CSF drainage in aged mice was associated with improved learning and memory performance.^8, 70–74^ The cholinergic system has also been associated with hydrocephalus in several prior studies, which supports a ChAT-dependent CSF circulation through the brain and spinal cord. In our PHH model, we observed significant differences in AuNP+/ChAT+ cell groupings between control and PHH rodents, where there were significant decreases in AuNP delivery to ChAT cell groupings. This suggests that disruptions of fluid dynamics within the ventricles may affect intracerebral CSF delivery, with further implications for cognitive dysfunction in hydrocephalus.^75, 76^ ACh and/or ChAT mediated CSF delivery to the brain may underly CSF pathophysiology during development and in neurodegenerative disease and deserves further study.

In our study, we also identify CSF flow across the ependyma of the lateral and third ventricles into neurogenic niches, including the subventricular zone, the hippocampus, and the olfactory bulb (via the rostral migratory stream). The differential and regional transependymal movement of large CSF solutes across ependymal walls and the leptomeninges with proximity to the hippocampus, pineal gland, hypothalamus, and corpus callosum may implicate a biological vulnerability of these structures to CSF abnormalities. Alternatively, the movement of CSF and CSF solutes into these areas may have a role in communication between the CSF and cells within the brain parenchyma. In development, it is possible that this transependymal movement of CSF represents a key mechanism by which growth factors reach the hippocampus and SVZ.^77–81^ This interpretation is consistent with previous studies that implicate growth factors, neurotransmitters, and other molecules pertinent in various signaling pathways diffuse across the ventricle wall to interact with type A, B2, and C cells in the SVZ.^79–81^ Changes in CSF composition and circulation dynamics in the setting of IVH-PHH may disrupt the interaction of these cells with transependymally transported molecules critical for development, with implications for ventricular zone disruption, neural stem cell loss, and eventual neurodevelopmental outcomes.

In this study, we used two tracers, 1.9 nm and 15 AuNPs, which both are coated with PEG and have a negative zeta potential. Previous have shown nanoparticle trophism is largely depending on surface chemistry but have not implicated a preferential interaction of PEG-coated NPs with specific cell types.^82, 83^ Accordingly, it cannot be ruled out that the 1.9 nm AuNP distribution patterns to cholinergic neurons is affected by a trophism for the AuNPs to these neurons, however this is unlikely as similar patterns of CSF tracer distribution were observed with other small molecule tracers with variable surface chemistry (Extended Data Figure 4). Furthermore, while we identified ChAT expression to be highly similar to 1.9 nm AuNP distribution, there may be other proteins that also have highly similar expression patterns, in addition to ChAT. Future studies using unbiased localization methods should be conducted to identify additional candidate markers.

This work describes, to our knowledge, the first CNS-wide map of anatomic brain and spine regions which preferentially see and handle CSF. We demonstrate the advantages to using such a high-resolution, global imaging modality to identify novel regions of interest for future CSF studies, highlighting the importance of the basal aspects of the brain to our understanding of CSF influx and circulation. These results provide a framework for the study of CSF-brain and CSF-spine interactions based not only on location, but also solute size. This work has implications for CSF circulation in the modulation of neurogenesis during neurodevelopment, CSF circulation pathologies including hydrocephalus, and pathological changes in aging with altered clearance of large proteins.

## Methods

### Animals (Rodents).

All experiments were approved by the institution’s Animal Care and Use Committee (protocol #19-0905). Sprague Dawley Rats (Charles River Laboratories, Wilmington, MA) were used in all experiments. To create the control and intraventricular hemorrhage conditions, we modified our previously established rodent model of PHH.^59^ Anesthetized (isoflurane 2.5% induction and 1.5% maintenance) P4 rats were fixed in a stereotaxic frame, a 2.5 mm midline incision was made and a 0.3cc syringe with a 30-gauge needle inserted into the right lateral ventricle (1.5 mm lateral, 0.4 mm anterior, and 2.0 mm deep). A small volume (20 µL) of artificial CSF (aCSF) (Tocris Bioscience, Bristol, UK) or hemoglobin (150 mg/ml diluted in sCSF) (EMD Millipore Corp, Burlington, Massachusetts) was injected at a rate of 8000 nL/min using a micro-infusion pump (World Precision Instruments, Sarasota, FL) to create the control and IVH-PHH conditions respectively. The needle was left in for 5 minutes post injection to prevent backflow. Incision was closed with 6-0 Ethilon suture (Ethicon Inc, Raritan, NJ). Rodents recovered from anesthesia and were returned to their cage with the mother.

### MRI quantification of ventricle volume

MRI was obtained 72 hours after intraventricular Hb or aCSF injection and images were acquired using a 4.7T Varian MRI scanner (Varian Inc., Palo Alto, California) with T2-weighted fast spin echo sequences (repetition time 3000/echo time 27.50 mS, 3 averages, field of view was 18.0 mm x 18.0 mm, matrix 128 x 128, 24 axial slices, and 0.50 mm thick). Rodents were anesthetized using isoflurane (2.5% induction, 1.5% maintenance), placed in a plastic holder and kept warm using a heated air blower. Lateral ventricles were segmented and quantified using ITK-SNAP software.

### AuNP size and zeta potential confirmation

The hydrodynamic size and morphology of 1.9 and 15 nm AuNPs (1102 and 1115, Nanoprobes, Yaphank, NY) were confirmed using transmission electron microscopy (TEM) with JEOL JEM-1400Plus Transmission Electron Microscope (JEOL, Peabody, Massachusetts) operating at 120 kV. A 2 µL sample was placed on an ultrathin lacey carbon grid (Formvar/Carbon 200 mesh, Copper, Ted Pella Inc., Redding, California) and allowed to sit for 5 min before removal of the droplet via wicking with a Chemwipe and vacuum drying.

Hydrodynamic diameter and zeta potential of the suspensions were determined with the Malvern Zetasizer Nano-ZS ZEN 3600 (Malvern, Malvern, United Kingdom) at 25 °C. Particle measurements were performed in a 1 cm path-length quartz cuvette and a folded capillary zeta cell (Malvern Instruments Ltd, Malvern, United Kingdom), respectively. Samples were highly diluted (c < 0.1 wt%) to prevent multiple scattering. A triplicate of the sample was diluted in DPBS for dynamic light scattering (DLS) and deionized water (diH2O) for zeta potential measurements.

Z-average size and polydispersity index (PDI) of the AuNPs were obtained with an average of 12 runs. TEM was performed to validate morphology and size of the AuNPs.

### AuNP tracer injection

1.9 and 15 nm AuNPs (1102 and 1115, Nanoprobes, Yaphank, NY) were constituted in aCSF at a concentration of 200 mg/mL. 72 hours after the initial Hb or aCSF injection, anesthetized P7 rats underwent intraventricular injection of 20 µl AuNPs as per protocol above with the following coordinates from bregma (1.7 mm lateral, 0.5 mm anterior, and 2.0 mm deep). Rodents were sacrificed at 10 minutes and 4 hours after the 1.9nm and 15 nm gold injections, respectively. For clearance studies, rodents were sacrificed 4 and 72 hours after the 1.9 nm and 15 nm gold injections respectively. A subset of rodents that were all used for the clearance and timepoint studies for 15 nm gold nanoparticle histology (Figure 5) underwent gold nanoparticle injection alone at P7, without preceding aCSF or Hb intraventricular injections.

### XRM acquisition and analysis

Following sacrifice, rodents were perfusion-fixed with 2x 10 mL 4% paraformaldehyde at 4° C and 10 mL ice-cold PBS and then placed in 4% paraformaldehyde at 4° C for transport to the imaging facility, embedded in 2% agarose, and were imaged within 24-72 hours of sacrifice using a Zeiss Versa 520 X-ray microscope (Carl Zeiss Imaging, White Plains, NY) using a 0.4x flat panel detector. X-ray source was tuned to 50 KV at 4W, to optimally excite gold particles. For samples requiring larger fields of view, we used the Versa’s wide or vertical stitch functions, as needed. A total of 1601 projections were acquired, reconstructed, and tomograms were visualized in either Zeiss XM3DViewer 1.2.8 (Carl Zeiss Imaging, White Plains, NY) or in Dragonfly 2020.1 (Object Research Systems, Montreal, Quebec). Processed images were systematically reviewed, and three images from each structure of interest were selected.

For standardization, images were numbered by their location relative to bregma, so the location of the selected images was consistent across all rodents. Three areas were selected in each anatomic location, and the mean gray intensity of these areas was calculated using ImageJ.

### Skull stripping and semi-automatic segmentation

XRM images were registered to the Duke Wistar rat atlas in-house “4dfp” tools (http://4dfp.readthedocs.io). Images were downsampled to 40 µm and transformation was computed to register images to the p8 average gradient echo image of the Duke Wistar atlas. This transformation was then applied to the full-resolution images. The co-registered atlas image (included only brain) was then used as a mask such that only the intra-cranial contents of each image remained. After skull-stripping, regions of intraparenchymal CSF were semi-automatically segmented to reveal the same patterns of CSF distribution (Figure 3).

### AuNP localization using light microscopy

After XRM image acquisition, the rodent was removed from the agarose and left in 4% PFA overnight. The brains were harvested and washed 2x in PBS for 1 hour each, then immersed in 30% and 50% ethanol for 30 minutes each, before being transferred to 70% ethanol for 24 to 72 hours at 4 degrees C prior to xylene and paraffin processing. The brains were embedded in paraffin, and 10 µm thick slices were sectioned in either the coronal or sagittal planes using a microtome to examine distribution of AuNPs.

Following overnight incubation at 60 °C, sections were soaked in xylene for 20 minutes and mounted with Permount mounting medium (#SP15-100, Thermo Fisher Scientific, Waltham, MA) for light microscopy localization of AuNPs.

To co-stain gold sections with cresyl violet, sections were hydrated through descending grades of alcohol (100%, 95%, 70%, 50%, 30%) to double distilled water (DDW) following xylene and immersed in FD Cresyl Violet Solution (#PS102-02, FD NeuroTechnologies, Columbia, MD) for 5 minutes. Sections were left in running DDW for 5 minutes then differentiated and rehydrated through ascending grades of alcohol (30%, 50%, 70%, 95%, 100%). Sections were then cleared in xylene for 20 minutes before mounting with Permount mounting medium (#SP15-100, Thermo Fisher Scientific, Waltham, MA).

### SEM imaging

Samples were incubated in a fixative containing 2.5% glutaraldehyde and 2% paraformaldehyde in 0.15 M cacodylate buffer, pH 7.4 overnight at 4°C. Post fixation, samples were rinsed in 0.15 M cacodylate buffer 3 times for 10 minutes each followed by a secondary fixation in 1% OsO4 in 0.15 M cacodylate buffer for 60 minutes in the dark. The samples were then rinsed 3 times in ultrapure water for 10 minutes each and dehydrated in a graded ethanol series (30%, 50%, 70%, 90%, 100% x2) for 10 minutes each step. Once dehydrated, the samples were then loaded into a critical point drier (Leica EM CPD 300, Vienna, Austria) which was set to perform 12 CO2 exchanges at the slowest speed. Once dried, samples were mounted on aluminum stubs with carbon adhesive tabs and coated with 20nm of carbon (Leica ACE 600, Vienna, Austria). SEM images were acquired on a FIB-SEM platform (Helios 5 UX DualBeam, Thermo Fisher Scientific, Brno, Czech Republic) using SEM imaging mode at 5 kV and 0.1 nA with TLD and ICD detectors.

### Anatomical analyses

Anatomical analyses identifying the location of AuNP circulation were performed using Brain Maps 4.0^42^ and The Rat Brain in Stereotaxic Coordinates (Seventh _Edition)._43

### Zebrafish Injections

Zebrafish were raised under standard conditions following IACUC guidelines. A casper line was crossed to our mitfa^-/-^Tg(HuC:EGFP) line and embryos were screened for EGFP signal at 24 hours post fertilization (hpf) and kept at 28.5C. Glass capillaries (TW100F-4, World Precision Instruments, Sarasota, FL) were pulled to desired shape using a Sutter Inst. P-97 model Micropipette Puller (Sutter Instrument, Novato, CA). At 96hpf, the fish were anesthetized in accordance with IACUC guidelines with tricaine methanesulfonate. During anesthetization, the tip of the pulled capillary was broken, and the capillary was loaded with an 8mg/mL Rhodamine B isothiocyanate-Dextran 70kDa (R9379, Millipore Sigma, Burlington, MA) in 1x aCSF. The microneedled capillary was loaded into a micromanipulator and the larval fish were placed on an agarose injection plate (under design secrecy) and oriented dorsal side up. The needle targeted the dorsal point along the midsagittal line along the border of the tectum and the cerebellum (targets the diencephalic ventricle). 1nL of the rhodamine solution was injected into the ventricles and the fish were returned to regular egg water. After 90 minutes, the fish were imaged using the VAST Bioimager (Union Biometrica, Holliston, MA) and a fluorescent confocal microscope (Carl Zeiss Imaging, White Plains, NY). ACSF used in zebrafish experiments contained: 125.5 mM NaCl, 20 mM NaHCO3, 2.4 mM KCl, 0.5 mM KH2PO4, 1.1 mM CaCl2•2H20, 0.85 mM MgCl2•6H20, 0.5 Na2SO4, 5 mM glucose; pH 7.4.

### In vivo gadolinium-enhanced MRI

P7 rodents were anesthetized with isoflurane and 0.5 µl of gadoterate meglumine solution (5mmol/mL Dotarem, Guerbet, Princeton, NJ) was injected into the right lateral ventricle using stereotaxic coordinates from the bregma (1.7mm lateral, 0.5mm anterior, 2.0mm deep). Rats were then placed in the Bruker 9.4T MRI (Bruker, Billerica, MA) and imaged 30 minutes post injection using a Bruker CryoProbe coil (Bruker, Billerica, MA) and the following T1-weighted paremeters: repetition time 800/echo time 7.8195 mS, 4 averages, field of view 24.0 x 24.0 mm, matrix 256 x 256, 24 axial slices, 0.50mm thick, and 14 sagittal slices, 1.0mm thick. Images were analyzed using the ITK-Snap software. Due to the heavy T1 weighting, very high concentrations of tracer appear dark, while lower concentrations appear bright. Areas of the brain without tracer distribution appear gray.

### Immunohistochemistry

All immunohistochemistry was performed on rodents with no AuNP injection as AuNP’s block light precluding chromogen or flurophore-based imaging. P7 rodents were anesthetized using 2% isoflurane and euthanized with CO2 inhalation before perfusion with 10 mL 4% paraformaldehyde at 4° C and 10 mL ice-cold PBS. Brains were harvested and left in 4% PFA overnight before being washed 2x in PBS for 1 hour, then immersed in 30% and 50% ethanol for 30 minutes each, before being left in 70% ethanol for 72 hours at 4 degrees C before xylene and paraffin processing. The brains were embedded in paraffin, and 10 µm thick slices were sectioned in either the coronal and sagittal planes and left in the incubator at 60 degrees overnight. Sections were then soaked in xylene to remove paraffin, before hydration through descending grades of alcohol (100%, 95%, 70%, 50%, 30%) to double distilled water (DDW).

Antigen retrieval was performed using a pH 6.0 citrate buffer (Sigma-Aldrich, St. Louis, MO) by microwaving in a 1:1000 solution in DDW for 30 minutes. Slides were cooled to room temperature, rinsed with PBS and blocked in 5% normal goat serum, 2.5% BSA, 0.5% TX-100 in PBS for one hour. Sections were incubated with appropriate dilutions of primary antibodies, 1:200 dilution of anti-ChAT antibody (#181023, Abcam, Cambridge, MA), 1:200 dilution of anti-NeuN antibody (#AB377, Sigma Aldrich, St. Louis, MO), 1:200 dilution of anti-CD31 antibody (#28364, Abcam, Cambridge, MA), 1:200 dilution of anti-S100β (#52642, Abcam, Cambridge, MA), 1:200 dilution of anti-GFAP (#7260, Abcam, Cambridge, MA) in PBS with 1% BSA and 0.5% TX-100 overnight at 4°C. Sections were then rinsed in PBS 6 times for 5 minutes each followed by incubation with goat anti-rabbit IgG Alexa Fluor 594 (#A32740, ThermoFisher Scientific, Waltham, MA) and goat anti-mouse IgG Alexa Fluor 488 (#A11008, ThermoFisher Scientific, Waltham, MA) diluted in PBS with 1% BSA, 0.5% TX-100 for 90 minutes at RT in a 1:2000 dilution. Sections were then rinsed in PBS 5 6 times 5 minutes each at room temperature followed by incubation with DAPI reagent in a 1:500 dilution. After 5 minutes in DAPI, sections were washed in PBS 3 times for 5 minutes and mounted with ProLong Gold Antifade Mountant (#P36930, Thermo Fisher Scientific, Waltham, MA). Histological evaluation was performed single blinded, by a trained observer without knowledge of the treatment.

### Meningeal wholemounts

Following AuNP injection, rodents were perfused with 10 mL 4% paraformaldehyde at 4° C and 10 mL ice-cold PBS. The cranium and spinal column were removed from the carcass. Curved micro scissors were used to carefully excise the brain and spinal cord, making sure to keep the dura and arachnoid overlaying the nervous tissues intact. The brain and spinal cord and overlying meninges were left in 4% PFA overnight at 4 degrees. The following day, the tissue was placed in a petri dish with ice-cold PBS. Under a dissecting microscope, the brain and spinal cord were first severed at the brainstem before the spinal cord was cut down its length (axially). The dorsal half of the spinal cord was secured with forceps and its adherent dura, which appeared as a loose-hanging translucent layer, was gently peeled away and mounted on a glass slide. Next, the dorsal leptomeninges were gently peeled away from the underlying parenchyma. This was repeated on the ventral half of the spinal cord. The cranial dura and leptomeninges were removed by adapting a previously reported protocol (Supplementary Methods).^84^ All wholemounts were transferred onto a glass slide, air dried, and mounted with Permount mounting medium (#SP15-100, Thermo Fisher Scientific, Waltham, MA).

### Statistical analysis

Rodents were randomly assigned to receive CSF or Hb injection to create the control and PHH groups. Statistical methods were not used to recalculate or predetermine sample sizes. Experimenters were blinded to the identity of experimental groups. Associations between two continuous variables were assessed using an unpaired T-test, and associations between more than two continuous variables were assessed using a one-way ANOVA. Mean integrated density for each anatomic location was compared to the other anatomic locations, across three different rodents. All tests were 2-tailed, and p-values of less than 0.05 were considered statistically significant. All analyses were performed using Microsoft Office Excel (Version 16.36) or GraphPad Prism (Version 9.0.0)

## Abbreviations

CSF: cerebrospinal fluid
MCA: middle cerebral artery
CNS: central nervous system
PHH: post-hemorrhagic hydrocephalus
IVH: intraventricular hemorrhage
SAS: subarachnoid space
AuNP: gold nanoparticle
XRM: X-ray microtomography
aCSF: artificial CSF
SSS: superior sagittal sinus
H: habenula
3 CN nucleus: oculomotor nucleus
DTg: dorsal tegmentum
IX: lamina IX of the ventral horn of the spinal cord
DPF: days post-fertilization
ChAT: choline acetyltransferase
ACh: acetylcholine
SVZ: subventricular zone
RMS: rostral migratory stream
AQ-4: aquaporin-4
AAV9: adeno-associated virus 9
CP: choroid plexus
3V: third ventricle

## Acknowledgements

We would like to thank Drs. Steven Brody and Jeffrey Neil for editing the manuscript and for their valuable comments and multiple discussions of this work. This work was funded by the National Institutes of Health (R01 NS110793 to J.M.S., P50HD103525 to A.Z.S., OD021694 to J.A.F), K12 Neurosurgeon Research Career Development Program (J.M.S), the Hydrocephalus Association (J.M.S), and by the McDonnell Center for Systems Neuroscience at Washington University in St. Louis. X-ray microtomographic imaging was performed using a Zeiss Xradia 520 X-ray Microscope (OD021694 to J.A.F), at the Washington University Center for Cellular Imaging (WUCCI), which is supported by Washington University School of Medicine, The Children’s Discovery Institute of Washington University and St. Louis Children’s Hospital (CDI-CORE-2015-505 and CDI-CORE-2019-813) (J.A.F.) and the Foundation for Barnes-Jewish Hospital (3770) (J.A.F).; The MRI studies presented in this work were supported by the Department of Radiology and were carried out in the Small Animal MR Facility of the Mallinckrodt Institute of Radiology at Washington University (S10OD026913-01 to J.D.Q). Figures were created in part with BioRender.com.

---

**Extended Data Figure 1.**
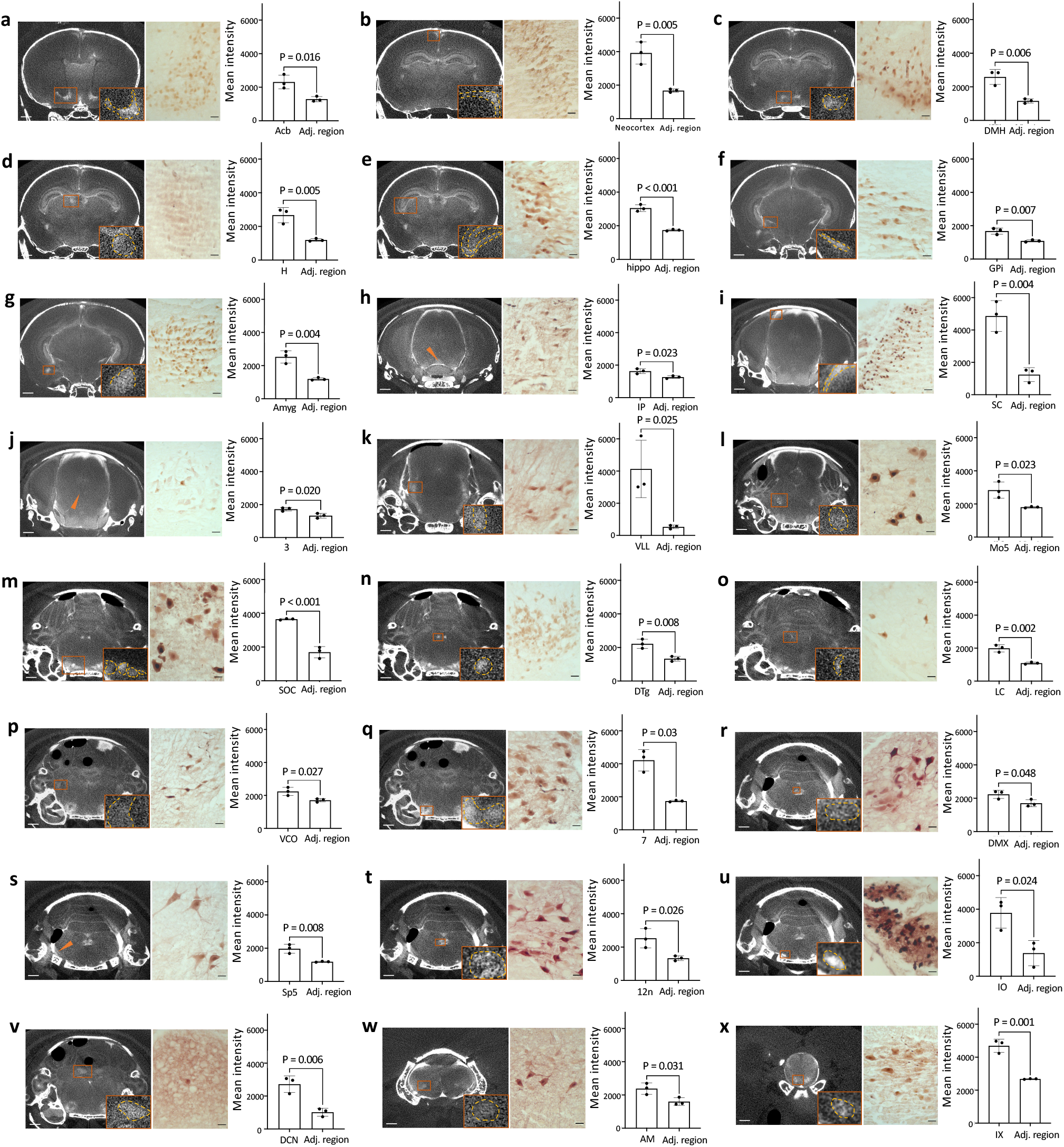
Neuron and stem cell distribution of intraventricularly injected small CSF solutes in the brain and spinal cord parenchyma. **a-w,** X-ray microtomography (XRM) image, histology and quantification of mean intensity of 1.9-nm gold nanoparticles (AuNPs) showing 1.9-nm AuNP distribution in the nucleus accumbens (acb) (a), neocortex (b), dorsal medial hypothalamus (DMH) (c), habenula (H) (d), hippocampus (hippo) (e), globus pallidus internus (GPi) (f), amygdala (Amyg) (g), interpeduncular nucleus (IP) (h), superior colliculus (SC) (i), oculomotor nucleus (3) (j), ventral lateral lemniscus (VLL) (k), motor nucleus of the trigeminal nerve (Mo5) (l), periolivary region (m), dorsal tegmentum (DTg) (n), locus coeruleus (LC) (o), ventral cochlear nucleus (VCO) (p), facial nerve nucleus (7) (q), dorsal motor nucleus of the vagus nerve (DMX) (r), spinal trigeminal nucleus (Sp5) (s), hypoglossal nucleus (12n) (t), inferior olivary nucleus (IO) (u), deep cerebellar nucleus (DCN) (v), nucleus ambiguus (AM) (w), lamina IX of the spinal cord (IX) (x). All regions above were significantly enhanced on the XRM images compared to surrounding parenchyma with visible 1.9-nm AuNPs on histological analysis. Data are mean ± s.e.m., n = 3 rodents, unpaired two-tailed t-test. XRM scalebar = 1 mm, histology scalebars = 25 µm.

**Extended Data Figure 2.**
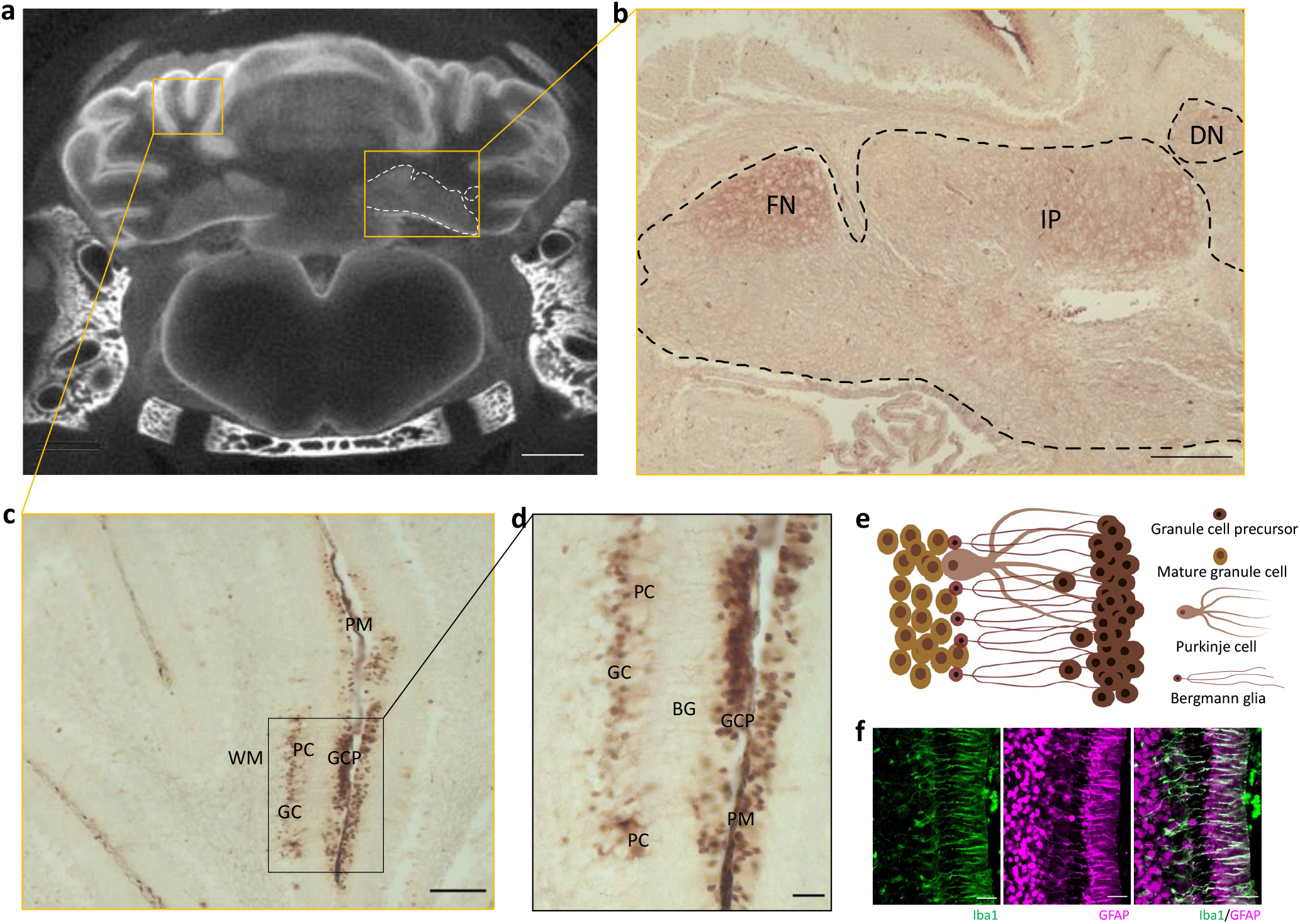
Intraventricularly injected CSF solutes circulate in and around the cerebellum. **a,** Gold nanoparticle enhanced X-ray microtomography (AuNP-XRM) showing distribution of 1.9-nm AuNPs in the cerebellum 10 minutes after injection into the right lateral ventricle of P7 aCSF control rodents. 1.9-nm AuNP distribution in the deep cerebellar nuclei is indicated with a white dashed line. scalebar = 1 mm. **b,** Representative histology showing 1.9-nm AuNP distribution in deep cerebellar nuclei (black dashed line), including the fastigial nucleus (FN), interposed nucleus (IPN), and dentate nucleus (DN). scalebar = 200 µm. **c-d,** Representative histology showing 1.9-nm AuNP distribution in the leptomeninges in the folia (PM, pia mater), granule cell precursors (GCP) and granule cell layer (GC). 1.9-nm AuNPs were also observed in Purkinje cells (PC) and fibers of Bergman glia (BG). C scalebar = 100 µm, d scalebar = 25 µm. **e,** Schematic showing organization of cerebellar cells observed with 1.9-nm AuNP staining in c and d. **f,** Ionized calcium binding adaptor molecule 1 (Iba1) (green) and glial fibrillary acidic protein (GFAP) (magenta) immunohistochemistry showing organization of BG and other glia in the cerebellum. scalebars = 50 µm. All data representative from 3 rodents.

**Extended Data Figure 3.**
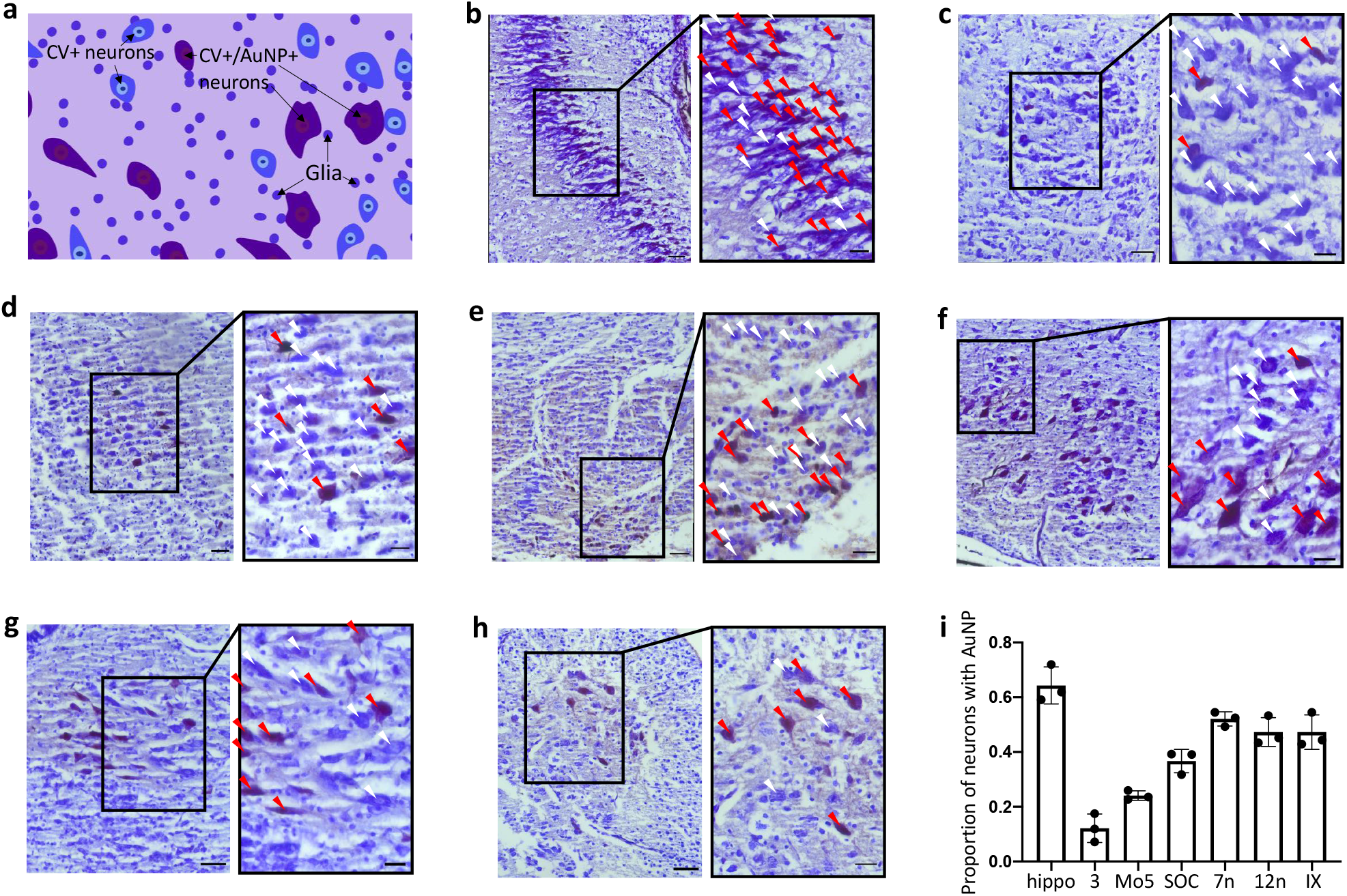
1.9-nm gold nanoparticle (AuNP) circulation in brain and spinal cord nuclei is not uniform across all neurons. **a,** Representative schematic showing distribution of neurons that are stained with cresyl violet (CV) only (CV+), CV and 1.9-nm AuNPs (AuNP+/CV+), and CV-stained glia. CV+ neurons are blue while CV+/gold+ neurons are violet. **b-h,** Representative micrograph showing CV co-staining with 1.9-nm AuNPs on a section of the hippocampus (b), oculomotor nucleus (c), motor nucleus of the trigeminal nerve (d), periolivary region (e), facial nucleus (f), hypoglossal nucleus (g), and lamina IX of the spinal cord (h) 10 minutes after 1.9-nm AuNP injection into the right lateral ventricle of P7 aCSF control rodents. Neurons stained with CV only are labeled with white arrowheads while neurons stained with CV and 1.9-nm AuNPs are labeled with red arrowheads. i, 1.9-nm AuNPs were observed in 64% of the CV+ neurons in the hippocampus, 12% of the CV+ neurons in the oculomotor nucleus, 25% of the CV+ neurons in the motor nucleus of the trigeminal nerve, 37% of the CV+ neurons in the periolivary region, 52% of the CV+ neurons in the facial nucleus, 47% of the CV+ neurons in the hypoglossal nucleus, and 47% of the CV+ neurons in lamina IX. Data are mean ± s.e.m., n = 3 rodents. scalebars = 50 µm, inset scalebars = 25 µm.

**Extended Data Figure 4.**
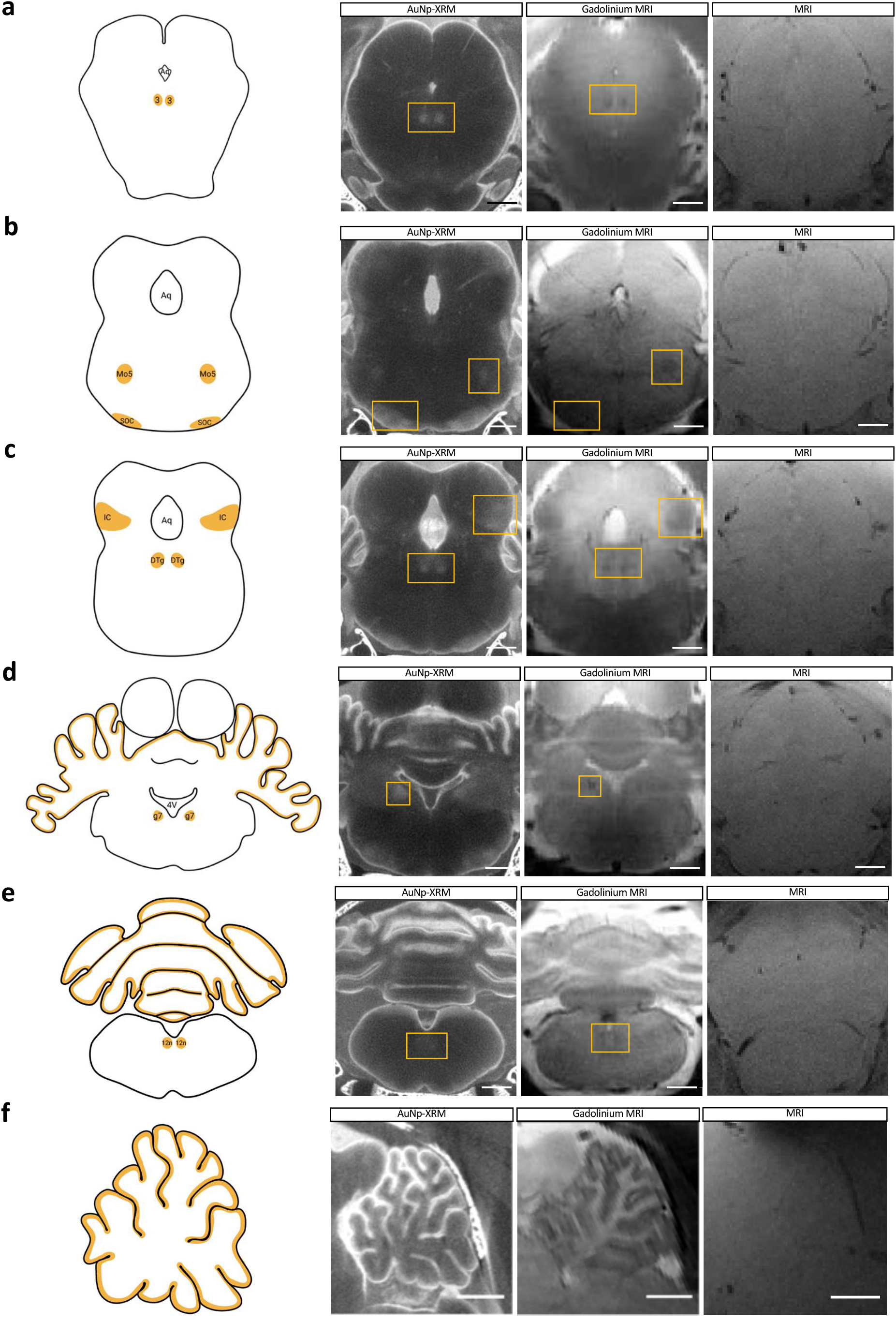
Gold nanoparticle-enhanced X-ray microtomography (AuNP-XRM) demonstrates similar patterns of intraparenchymal CSF tracer distribution when compared to in-vivo gadolinium-enhanced MRI studies. **a-f,** AuNP-XRM (left) performed 10 minutes after 1.9-nm AuNPs were injected into the right lateral ventricle of P7 aCSF control rodents and in-vivo gadolinium enhanced MRI (middle) performed 30 minutes after gadolinium injection into the right lateral ventricle of P7 aCSF control rodents show similar patterns of tracer distribution in the oculomotor nucleus (3) (a), motor nucleus of the trigeminal nerve (Mo5) (b), periolivary region (POR) (b), inferior colliculus (IC) (c), dorsal tegmental nucleus (DTg) (c), genu of the facial nerve (g7) (d), cerebellum (d, e, f),and hypoglossal nucleus (12n) (e). Schematics showing locations of cell groupings and nuclei with tracer distribution are provided for reference (left). MRIs performed without gadolinium contrast agent are shown for comparison (right). scalebar = 1 mm. All data are representative of 3 rodents.

**Extended Data Figure 5.**
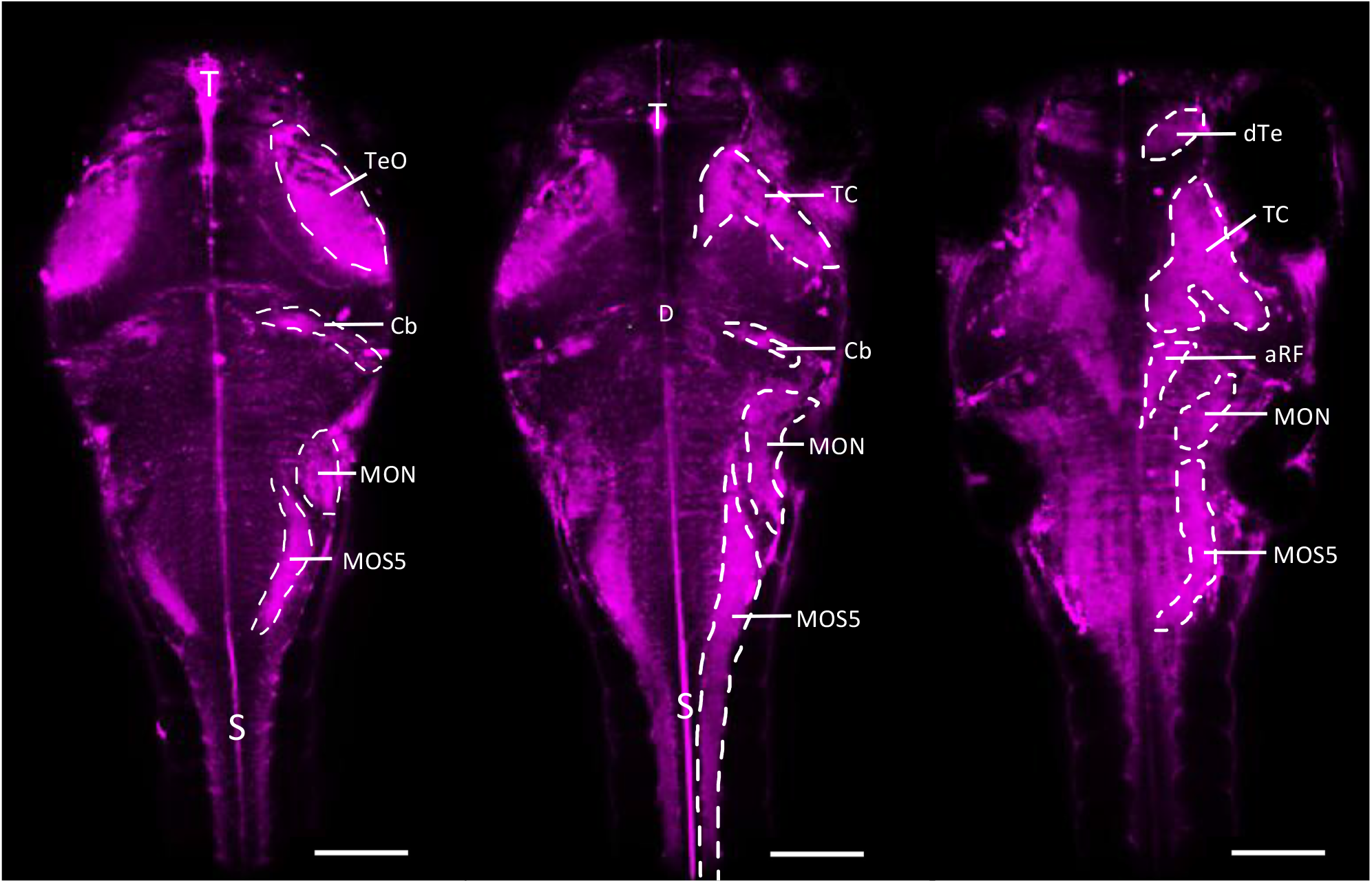
Fluorescent CSF tracers preferentially distribute to neuronally-rich populations in zebrafish, with similar patterns to gold nanoparticle enhanced X-ray microtomography (AuNP-XRM) in rodents. Dextran 70 kDa (magenta) distribution through the zebrafish brain and spinal cord 90 minutes post intraventricular injection in a 96 hpf zebrafish. Dextran is observed in the CSF spaces of the telencephalic ventricle (T), diencephalic ventricle (D), spinal canal (S), and areas of the parenchyma including the optic tectum (TeO), tectum (TC), cerebellum (Cb), medial octavalis nucleus (MON), inferior dorsal medulla oblongata stripe 5 (MOS5), dorsal telencephalon (dTe), and anterior reticular formation (aRF). scalebar = 1 mm. All data are representative of 3 zebrafish.

**Extended Data Figure 6.**
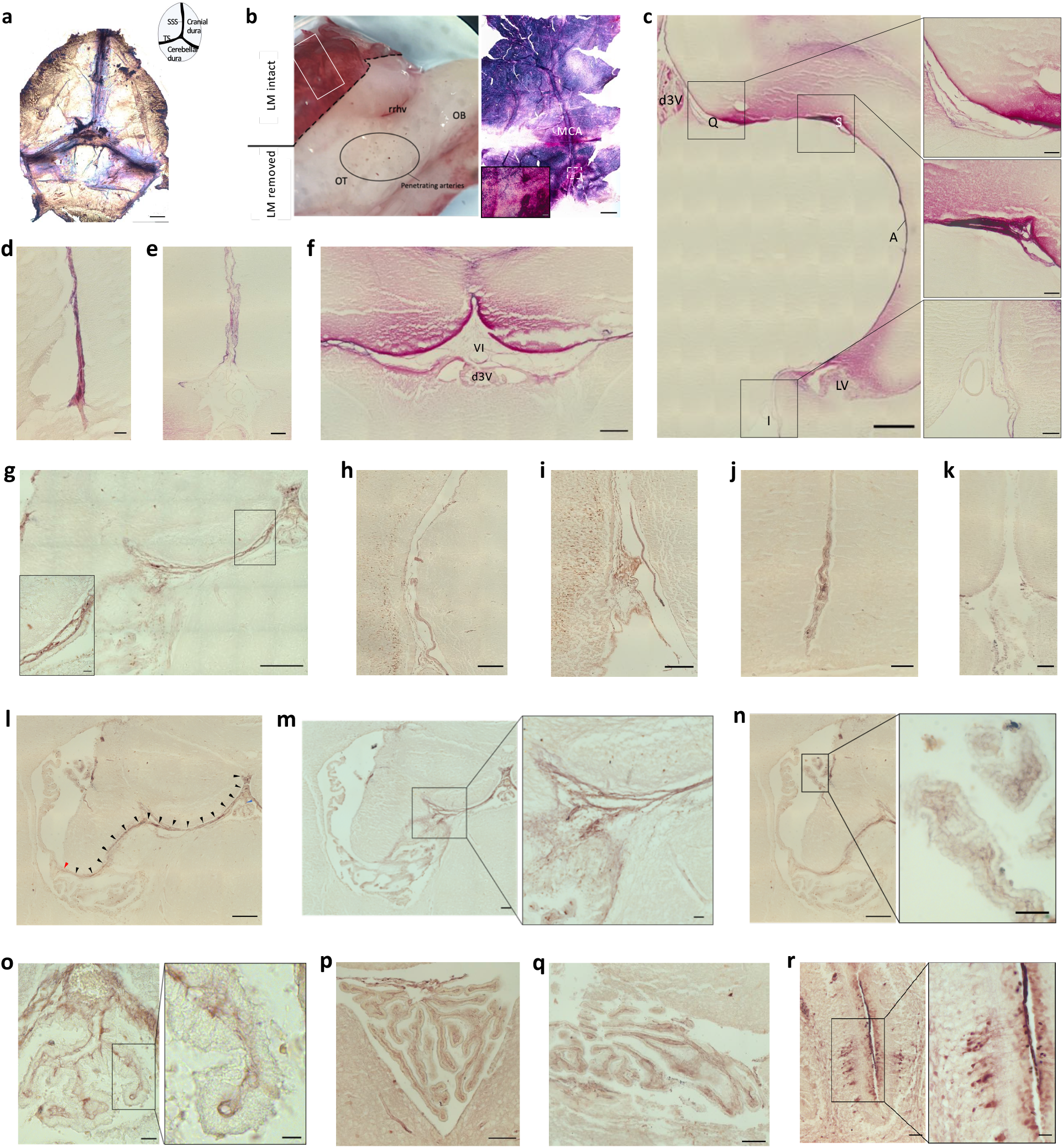
Intraventricularly injected large and small CSF solutes circulate in the leptomeninges of the brain. **a,** Representative image of 15-nm gold nanoparticle (AuNP) distribution through the dura 4 hours after intraventricular injection in P7 rodents. Inset, schematic demonstrating anatomy of whole-mount dissection of the rat brain dura is shown. SSS, superior sagittal sinus; TS, transverse sinus. scalebar = 1 mm. **b,** Dissection microscope image of harvested P7 rat brain 4 hours after 15-nm AuNP injection with the leptomeninges (LM) peeled back from the base of the brain. Solid white box indicates the location that the leptomeningeal wholemount (right) showing the middle cerebral artery (MCA) is taken from. Inset, higher magnification image of the MCA on the LM wholemount. olfactory tubercle; rrhv, rostral rhinal vein; OB, olfactory bulb. scalebar = 1 mm, inset scalebar = 50 µm **c,** Representative histology showing 15-nm AuNP circulation through the dorsal third ventricle (d3V), quadrigeminal cistern (Q), subarachnoid cistern (S), ambient cistern (A), interpeduncular cistern (I), and lateral ventricle (LV) 4 hours after intraventricular injection in P7 rodents. **d-f,** The longitudinal fissure (d) rhinal fissure (e), and velum interpositum (f) 4 hours after intraventricular 15-nm AuNP injection at P7. VI, velum interpositum; d3V, dorsal third ventricle. c scalebar = 500 µm, inset scalebars = 75 µm, d scalebar = 50 µm, e scalebar = 100 µm, f scalebar = 250 µm. **g-m,** Representative histology showing 1.9-nm AuNP circulation through the quadrigeminal cistern (g), ambient cistern (h), interpeduncular cistern (i), longitudinal fissure (j), rhinal fissure (k), pia mater of the tela choroidea (l) (black arrowheads), entry of the tela choroidea pia mater into the right lateral ventricle choroid plexus (m) (red arrowhead, m, inset m), and third ventricle choroid plexus (m) 10 minutes after intraventricular injection in P7 aCSF control rodents. g scalebar = 250 µm, g inset scalebar = 25 µm, h-i scalebar = 250 µm, j scalebar = 100 µm, k scalebar = 100 µm, l scalebar = 250 µm, m scalebar = 100 µm, inset scalebar = 50 µm. **n-q,** Representative histology of 1.9-nm AuNP circulation through the choroid plexus of the right lateral ventricle (n), dorsal third ventricle (o), fourth ventricle (p), and lateral fourth ventricle (q) 10 minutes after intraventricular injection in P7 aCSF control rodents. n scalebar = 250 µm, n inset scalebar = 50 µm, o scalebar = 25 µm, o inset scalebar = 10 µm, p scalebar = 25 µm, q scalebar = 100 µm **r,** Representative histology of 1.9-nm AuNP circulation through the folia of the cerebellum 10 minutes after intraventricular injection in P7 aCSF control rodents. All data are representative of 3 rodents.

**Extended Data Figure 7.**
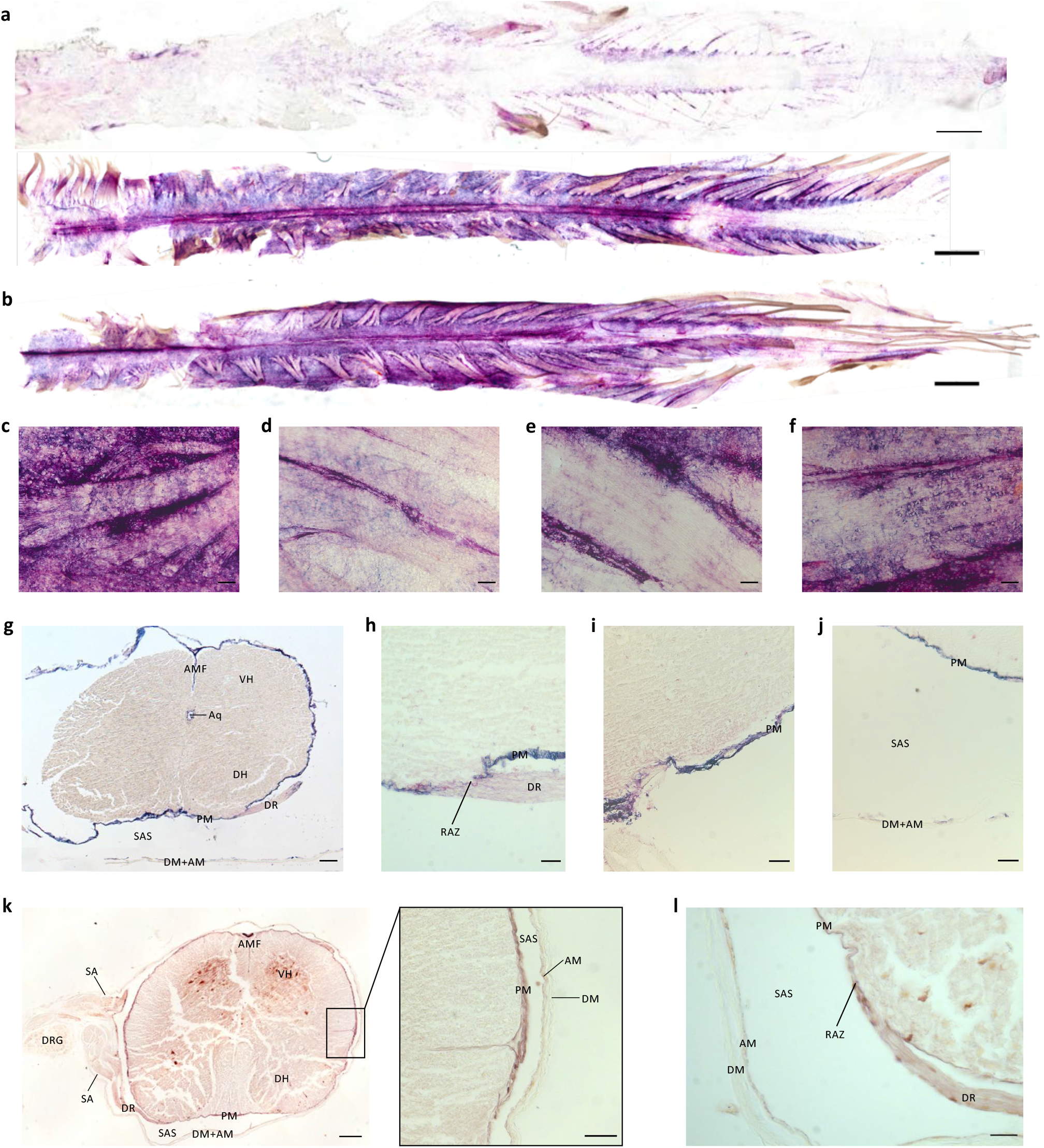
Intraventricularly 1.9-nm and 15-nm gold nanoparticles {AuNPs) circulate in the leptomeninges of the spinal cord. **a,** Representative wholemounts showing 15-nm AuNP distribution through the dorsal spinal cord dura (top) and dorsal spinal cord leptomeninges (bottom) 4 hours after intraventricular injection into the right lateral ventricle of P7 rodents. scalebar = 1 cm. **b,** Representative wholemount showing 15-nm AuNP distribution through the ventral spinal cord leptomeninges 4 hours after intraventricular injection into the right lateral ventricle of P7 rodents. scalebar = 1 cm. a-b, black arrows indicate 15-nm AuNPs forming the outlines of nerve root. **c-f,** High magnification histology of 15-nm gold distribution along the leptomeninges of spinal nerve roots as they leave the spinal cord. 15-nm AuNPs appeared to distribute primarily around the roots, with additional diffuse 15-nm AuNPs observed inside the roots. scalebars = 25 µm. **g-j,** Cross sectional histology showing 15-nm AuNP distribution in the spinal cord, leptomeninges and dura 4 hours after intraventricular injection into the right lateral ventricle of P7 rodents. The anterior median fissure (AMF), ventral horn (VH), aqueduct (Aq), dorsal horn (DH), dorsal root (DR), dura mater (DM), leptomeninges (LM) and root attachment zone (RAZ) are indicated. g scalebar = 1 mm, h-j scalebars = 25 µm. **k-l,** Cross sectional histology showing 1.9-nm AuNP distribution in the spinal cord, leptomeninges, and dura 10 minutes after intraventricular injection into the right lateral ventricle in P7 aCSF control rodents. The AMF, VH, subarachnoid angle (SA), dorsal root ganglion (DRG), DR, PM, subarachnoid space (SAS), DM and arachnoid mater (AM) are indicated. k scalebar = 1 mm, k inset scalebar = 25 µm, l scalebar = 25 µm. All data are representative of 3 rodents.

**Extended Data Figure 8.**
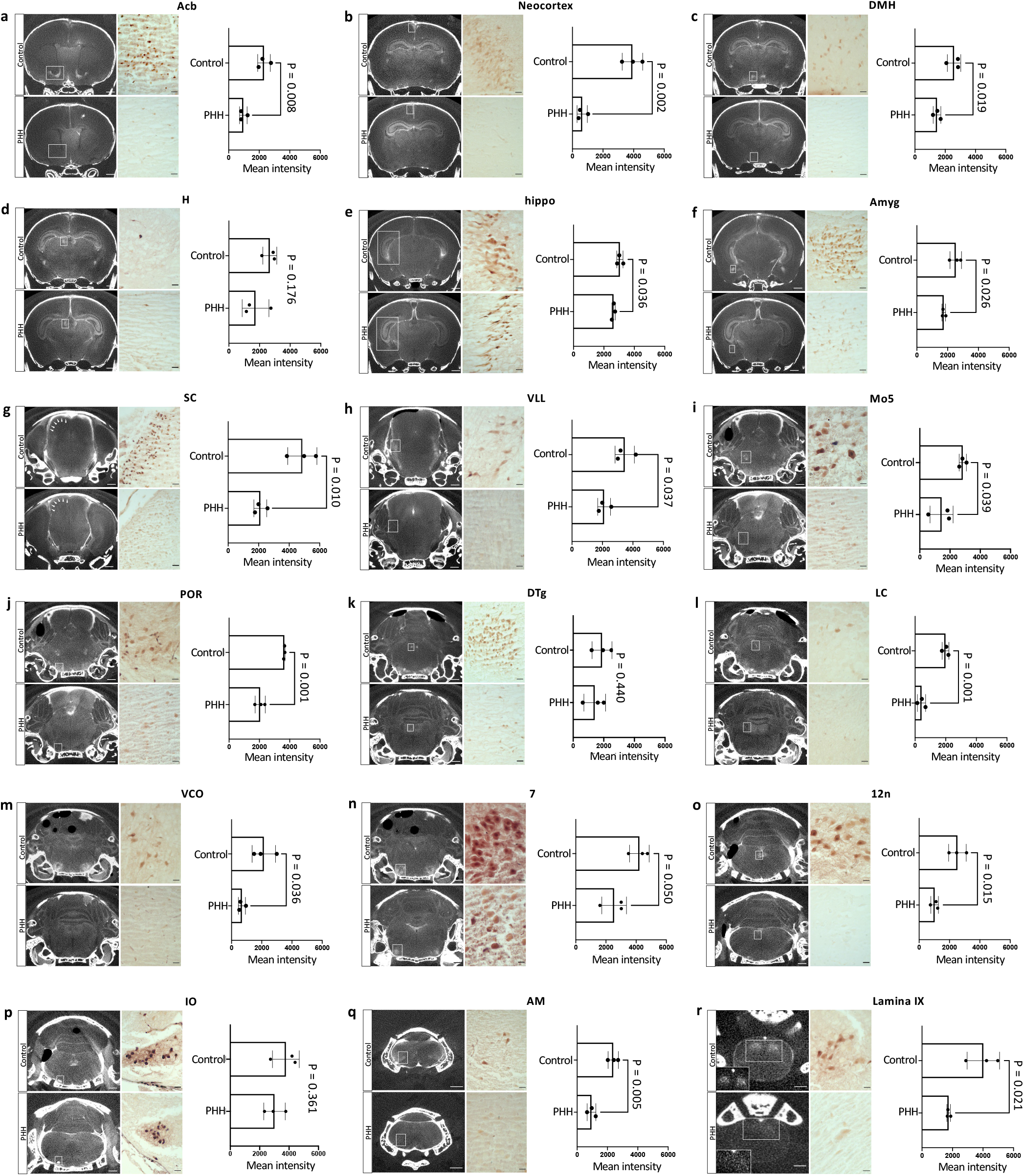
Altered targeted CSF distribution in the brain and spinal cord parenchyma in posthemorrhagic hydrocephalus (PHH). **a,** Representative coronal X-ray nanotomography (XRM), histology, and quantification showing altered 1.9-nm AuNP distribution in the nucleus accumbens (Acb) (a), neocortex (b), dorsal medial hypothalamus (DMH) (c), habenula (H) (d), hippocampus (hippo) (e), amygdala (Amyg) (f), superior colliculus (SC) (g), ventral lateral lemniscus (VLL) (h), motor nucleus of the trigeminal nerve (Mo5) (i), periolivary region (POR) (j), dorsal tegmental nucleus (DTg) (k), locus coeruleus (LC) (l), ventral cochlear nucleus (VCO) (m), facial nucleus (7) (n), hypoglossal nucleus (12n) (o), inferior olivary nucleus (IO) (p), nucleus ambiguus (AM) (q), lamina IX of the spinal cord (IX) (r) in P7 PHH rodents 10 minutes following intraventricular injection, compared to aCSF controls. scalebars = 1 mm, histology scalebars = 25 µm. Data are mean ± s.e.m., n = 3 rodents, unpaired two-tailed t-test.

**Extended Data Figure 9.**
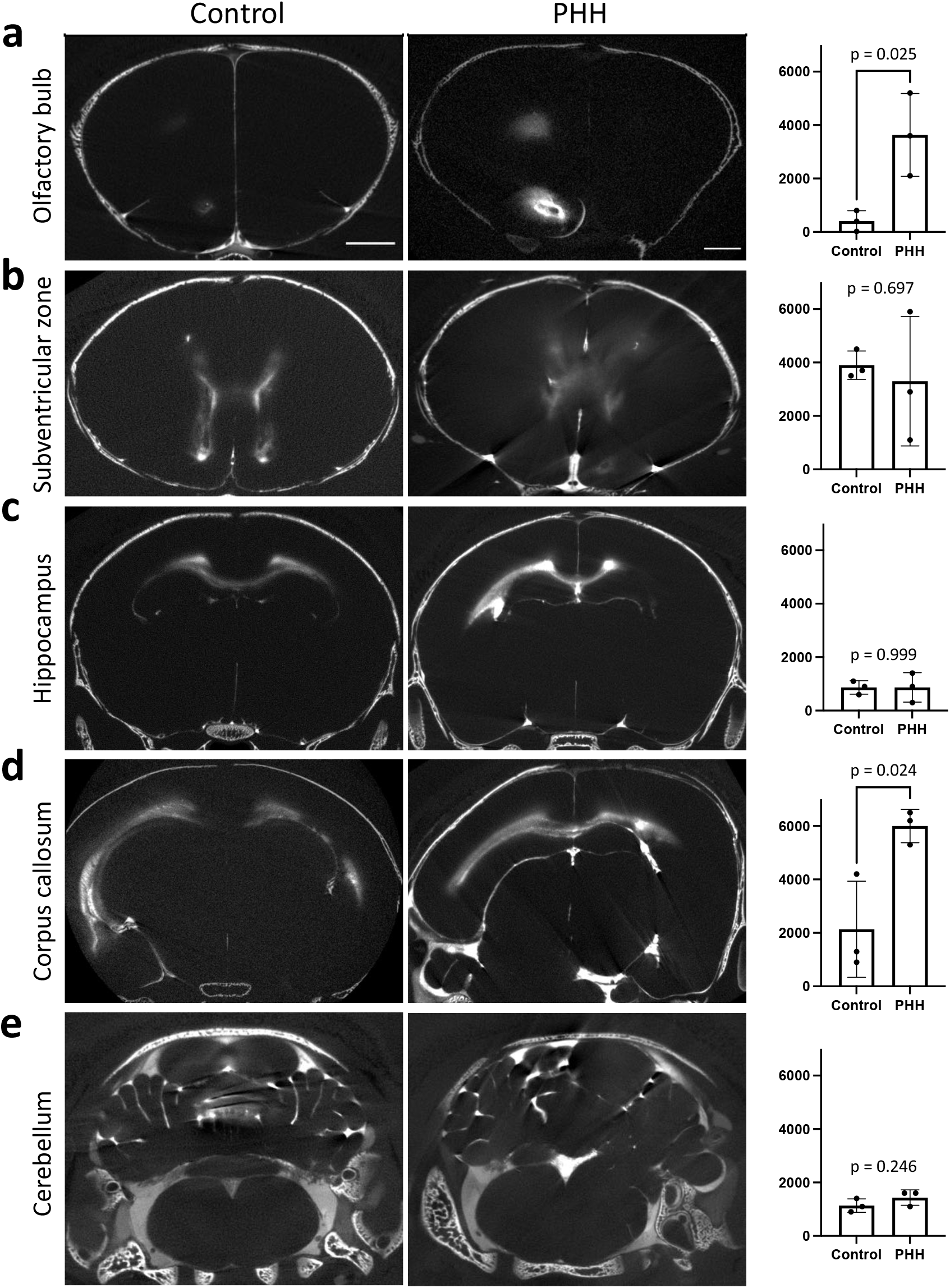
Altered distribution of 15-nm AuNPs in the brain and spinal cord parenchyma in posthemorrhagic hydrocephalus (PHH). **a-e,** Representative coronal XRM and quantification of 15-nm AuNP distribution to the olfactory bulb (a), subventricular zone (b), hippocampus (c), corpus callosum (d), and cerebellum (e). Data are mean ± s.e.m., n = 3 rodents, unpaired two-tailed t-test.

