## Supplementary Information for "Global cerebrospinal fluid circulation mapping using gold nanoparticle enhanced X-ray microtomography reveals region-specific brain and spinal cord CSF pathways"

| Region | ChAT+ cell bodies | Description | Reference |
| --- | --- | --- | --- |
| Nucleus accumbens | + | Observed on IF (Figure 4), shown in previous studies. | ^1,2^ |
| Neocortex | + | Observed on IF (Figure 4), shown in previous studies. | ^3–5^ |
| Hippocampus | + | Observed on IF (Figure 4), shown in previous studies. | ^6–9^ |
| Habenula | + | Observed on IF (Figure 4), shown in previous studies. | ^3,10^ |
| Globus pallidus internus | + | Observed on IF (Figure 4), shown in previous studies. | ^1^ |
| Dorsal medial hypothalamus | + | Observed on IF (Figure 4), shown in previous studies. | ^1^ |
| Amygdala | + | Observed on IF (Figure 4), shown in previous studies. | ^11–13^ |
| Oculomotor nucleus | + | Observed on IF (Figure 4), shown in previous studies. | ^1,14^ |
| Trochlear nucleus* | + | Shown in previous studies. | ^1,14^ |
| Superior colliculus | - | Not observed on IF. |  |
| Interpeduncular nucleus | + | Observed on IF (Figure 4), shown in previous studies. | ^10,15^ |
| Ventral lateral lemniscus | + | Observed on IF (Figure 4), shown in previous studies. | ^16,17^ |
| Periolivary region | + | Observed on IF (Figure 4), shown in previous studies. | ^18^ |
| Dorsal tegmentum | - | Not observed on IF. |  |
| Locus coeruleus | + | Observed on IF (Figure 4), shown in previous studies. | ^19,20^ |
| Ventral cochlear nucleus | + | Observed on IF (Figure 4), shown in previous studies. | ^21^ |
| Motor trigeminal nucleus | + | Observed on IF (Figure 4), shown in previous studies. | ^1,2^ |
| Spinal trigeminal nucleus | - | Not observed on IF. |  |
| Facial nerve nucleus | + | Observed on IF (Figure 4), shown in previous studies. | ^1,10^ |
| Nucleus ambiguus | + | Observed on IF (Figure 4), shown in previous studies. | ^1^ |
| Dorsal motor vagus nucleus | + | Observed on IF (Figure 4), shown in previous studies. | ^1^ |
| Hypoglossal nucleus | + | Observed on IF (Figure 4), shown in previous studies. | ^1,10^ |
| Inferior olivary nucleus | - | Not observed on IF. |  |
| Deep cerebellar nuclei | + | Observed on IF (Figure 4), shown in previous studies. | ^22–25^ |
| Lamina IX | + | Observed on IF (Figure 4), shown in previous studies. | ^1,26^ |

**Supplementary Table 1. ChAT positivity by nuclei and cell groupings with observed 1.9 nm AuNP distribution.**

| Region | Reference |
| --- | --- |
| Caudate/Putamen | ^1^ |
| Nucleus basalis of Meynert | ^27^ |
| Magnocellular preoptic nucleus | ^1^ |
| Olfactory tubercle | ^1^ |
| Islands of Calleja | ^1^ |
| Nucleus of the diagonal band | ^1^ |
| Substantia innominata | ^1^ |
| Medial septal nucleus | ^1^ |
| Septofimbrial nucleus | ^1^ |
| Bed nucleus of the stria terminalis | ^1^ |
| Septo-olfactory area | ^1^ |
| Edinger Westphal nucleus | ^1^ |
| Pedunculopontine tegmental nucleus | ^1^ |
| Superior cerebellar peduncle | ^1^ |
| Parabigeminal nucleus | ^1^ |
| Laterodorsal tegmental nucleus | ^1^ |
| Parabrachial nucleus | ^1^ |
| Abducens nucleus | ^1^ |

**Supplementary Table 2. Known ChAT-positive nuclei and cell groupings with no observed 1.9 nm AuNP distribution.**

| Region | Abbreviation |
| --- | --- |
| Oculomotor nucleus | 3 |
| Dorsal tegmentum | DTg |
| Habenula | H |
| Amygdala | Amyg |
| Central gray | CG |
| Motor nucleus of the trigeminal nerve | Mo5 |
| Periolivary region | POR |
| Deep cerebellar nuclei | DCN |
| Facial nerve root | 7n |
| Facial nerve nucleus | 7 |
| Nucleus ambiguus | AM |
| Inferior olivary nucleus | IO |
| Hypoglossal nucleus | 12n |
| Lamina IX of the spinal cord | IX |
| Interpeduncular nucleus | IP |
| Red nucleus | RN |
| Nucleus of Darkschewitsch | DK |
| Trochlear nucleus | 4 |
| Leptomeninges | LM |
| Caudal rhinal vein | crhv |
| Lateral ventricle | LV |
| Hippocampus | hippo |
| M1 of the middle cerebral artery | M1 |
| Lenticulostriate arteries | LSA |
| Ventral third ventricle | v3V |

**Supplementary Table 3. Neuroanatomical abbreviations for Figure 3.**

| Region | Abbreviation |
| --- | --- |
| Accumbens nucleus | Acb |
| Layers III and V of the motor cortex | III and V |
| Dorsomedial hypothalamus | DMH |
| Habenula | H |
| Hippocampus | hippo |
| Amygdala | Amyg |
| Ventral lateral lemniscus | VLL |
| Motor nucleus of the trigeminal nerve | Mo5 |
| Periolivary region | POR |
| Dorsal tegmentum | DTg |
| Locus coeruleus | LC |
| Ventral cochlear nucleus | VCO |
| Facial nucleus | 7 |
| Hypoglossal nucleus | 12n |
| Inferior olivary nucleus | IO |
| Deep cerebellar nucleus | DCN |
| Nucleus ambiguus | AM |
| Lamina IX of the spinal cord | IX |
| Superior colliculus | SC |
| Spinal trigeminal nucleus | Sp5 |
| Globus pallidus internus | GPi |
| Interpeduncular nucleus | IP |
| Oculomotor nucleus | 3 |
| Dorsal motor nucleus of the vagus nerve | DMX |
| Nucleus of Darkschewitsch | DK |
| Red nucleus | RN |
| Trochlear nucleus | 4 |
| Nucleus sagulum | SAG |

**Supplementary Table 4. Neuroanatomical abbreviations for Figure 8.**

**Supplementary methods**

*Meningeal wholemounts.* Following AuNp injection, animals were perfused with 10 mL 4% paraformaldehyde at 4º C and 10 mL ice-cold PBS and placed dorsal side up. A single midline cut was made from the nose to the tail and the skin retracted to expose the skull, muscle and underlying tissues. Spinal column excision was performed by making an incision in the region of the lower back/femurs 0.5 cm lateral to the column, and cutting the back musculature up towards the head, cutting through the hip joint, ribs, and shoulder joint to remove the arm and leg, with care taken to keep the central nervous system structures intact. This was repeated on the contralateral side. The viscera attached on the ventral side of the spinal column was cut to free the spinal column and head from the rest of the body. Using curved microsurgical scissors, with the tips pointing away from the brain, the posterior atlanto-occipital membrane was cut, exposing the foramen magnum. Sliding the curved scissors into the foramen magnum, the skull was cut counterclockwise, superior to the posttympanic hook and the zygomatic process of the frontal bone and around the anterior-most aspect of the frontal bone to remove the frontal, parietal, interparietal, and occipital bones and the adherent underlying dura in one piece. The curved scissors were then held parallel to the table, with tips pointing away from the spinal cord, and used to sever the pedicles of the vertebra to remove the vertebral bodies and expose the dorsal side of spinal cord. The scissors were rotated 90 degrees and used to cut the vertebral lamina and sever the dorsal roots as close to the cord as possible. The brain and spinal cord tissue were carefully separated from the remaining bone in one piece, taking care to keep the dura and arachnoid intact, and left in 4% PFA overnight at 4 degrees.

The following day, the tissue was placed in a petri dish with ice-cold PBS under a dissecting microscope. The brain and spinal cord were severed. Using curved forceps to gently secure the spinal cord, microsurgical scissors were used to cut the dura and leptomeninges along the length of both sides of the spinal cord. After orienting the spinal cord with the dorsal side facing up, the dura, which appeared as a loose, translucent layer, was gently peeled away from the leptomeninges and parenchyma in one piece. Next, the leptomeninges, which appeared as a spongy layer with distinctive AuNp presence, was gently peeled away from the underlying parenchyma. This was repeated on the ventral side. The cranial leptomeninges were removed in three pieces, one from each cerebral hemisphere, and one from the cerebellum. To remove the leptomeninges from the cerebral hemispheres, a midline incision was made in the leptomeninges on the base of the brain. Using curved forceps, the leptomeninges were carefully peeled from the incision moving up towards the longitudinal fissure and removed as one large piece. The cerebellar leptomeninges were removed by using the curved forceps to detach the leptomeninges from the superior colliculi and gently peeling tissue from the folia, moving down towards the foramen magnum.

Dura and pia-arachnoid wholemounts were transferred onto a glass slide, air dried, and mounted with Permount mounting medium (#SP15-100, Thermo Fisher Scientific, Waltham, MA) for imaging with light microscopy.

9.	GENSAT Project at Rockefeller University, Mouse Brain Atlas, Image Navigator. http://www.gensat.org/imagenavigator.jsp?imageID=29460.

10.	GENSAT Project at Rockefeller University, Mouse Brain Atlas, Image Navigator. http://www.gensat.org/imagenavigator.jsp?imageID=29464.

11.	Hellendall, R. P., Godfrey, D. A., Ross, C. D., Armstrong, D. M. & Price, J. L. The distribution of choline acetyltransferase in the rat amygdaloid complex and adjacent cortical areas, as determined by quantitative micro-assay and immunohistochemistry. *The Journal of comparative neurology* **249**, 486–498 (1986).

12.	Emson, P. C., Paxinos, G., le Gal La Salle, G., Ben-Ari, Y. & Silver, A. Choline acetyltransferase and acetylcholinesterase containing projections from the basal forebrain to the amygdaloid complex of the rat. *Brain research* **165**, 271–282 (1979).

13.	Ben-Ari, Y., Zigmond, R. E., Shute, C. C. D. & Lewis, P. R. Regional distribution of choline acetyltransferase and acetylcholinesterase within the amygdaloid complex and stria terminalis system. *Brain research* **120**, 435–445 (1977).

14.	GENSAT Project at Rockefeller University, Mouse Brain Atlas, Image Navigator. http://www.gensat.org/imagenavigator.jsp?imageID=29463.

26.	GENSAT Project at Rockefeller University, Mouse Brain Atlas, Image Navigator. http://www.gensat.org/imagenavigator.jsp?imageID=29467.

27.	Eckenstein’ And, F. & Sofroniew, M. v. IDENTIFICATION OF CENTRAL CHOLINERGIC NEURONS CONTAINING BOTH CHOLINE ACETYLTRANSFERASE AND ACETYLCHOLINESTERASE AND OF CENTRAL NEURONS CONTAINING ONLY ACETYLCHOLINESTERASE’. *Society for Neuroscience* **3**, 2286–2291 (1983).
